# Modeling phytoplankton-zooplankton interactions: opportunities for species richness and challenges for modern coexistence theory

**DOI:** 10.1101/2022.03.24.485680

**Authors:** Jurg W. Spaak, Peter B. Adler, Stephen P. Ellner

## Abstract

Many potential mechanisms can sustain biodiversity, but we know little about their relative importance. To compare multiple mechanisms, we modeled a two-trophic planktonic food-web based on mechanistic species interactions and empirically measured species traits. We simulated thousands of communities under realistic and altered trait distributions to assess the relative importance of three potential drivers of species richness: resource competition, predator-prey interactions, and trait trade-offs. Next, we computed niche and fitness differences of competing zooplankton to obtain a deeper understanding of how these mechanisms limit species richness. We found that predator-prey interactions were the most important driver of species richness and that fitness differences were a better predictor of species richness than niche differences. However, for many communities we could not apply modern coexistence theory to compute niche and fitness differences due to complications arising from trophic interactions. We therefore need to expand modern coexistence theory to investigate multi-trophic communities.

## Introduction

When Hutchinson coined the term “paradox of the plankton,” he was emphasizing that plankton all seem to belong to the same niche, at least to a naive observer (Hutchinson, 1959). But 60 years of research has produced a long list of the different limiting factors that could promote stable coexistence in plankton communities and the diversity of primary producers in general. For example, we expect trade-offs between resource affinities for light, nitrogen or phosphorus (Litchman *et al*., 2007; Litchman & Klausmeier, 2008; Edwards *et al*., 2011; Kraft *et al*., 2015) or for different wavelengths of light (Stomp *et al*., 2004; Spaak *et al*., 2021a). Moreover, we can expect some species to specialize on low resource conditions and others on high resource conditions, the “gleaner-opportunist” trade-off (Litchman & Klausmeier, 2001; Kiørboe *et al*., 2018). We expect a growth-defense trade-off, where some species invest in rapid growth while other species allocate resources to defense against predation (Finkel *et al*., 2010; Branco *et al*., 2020; Lind *et al*., 2013). We know that generalist consumers can coexist with species-specific consumers and, more generally, that predator-mediated effects can increase species richness (Janzen, 1970; Ehrlich *et al*., 2020; Olff & Ritchie, 1998; Bagchi *et al*., 2014; Becerra, 2015). Additionally, we have fluctuation dependent mechanisms, storage effects and relative non-linearities (Letten *et al*., 2018; Zepeda & Martorell, 2019; Litchman, 2003; Ellner *et al*., 2019), the interaction of evolution and fluctuations (Yamamichi *et al*., 2020), and internally generated fluctuations (Huisman *et al*., 2006). We now may have too many explanations for diversity rather than too few, as we know little about the relative importance of all these mechanisms (Shoemaker *et al*., 2019).

Modern coexistence theory is a general framework with which we can analyze these different mechanisms and compare them. Modern coexistence theory has successfully been applied to understand many facets of species richness by providing tools to quantify and compare multiple mechanisms affecting species richness (Barabás *et al*., 2018; Ellner *et al*., 2019; Chesson, 2000), and disentangling differences promoting coexistence, i.e. niche differences, form differences hampering coexistence, i.e. fitness differences (Carroll *et al*., 2011; Spaak & De Laender, 2020; Spaak *et al*., 2021c). For example, modern coexistence theory captures the gleaner-opportunist trade-off into the more general concept of relative non-linearity (Litchman & Klausmeier, 2001; Yamamichi *et al*., 2020; Letten *et al*., 2018). Storage effect, another well studied coexistence mechanism, captures the idea that covariance between good environments and low competition can benefit a species at low density (Chesson, 1994; Angert *et al*., 2009; Usinowicz *et al*., 2012; Shoemaker *et al*., 2020a; Usinowicz *et al*., 2017). Much attention has also been given to environments without fluctuations, where traits in plant or phytoplankton communities govern both, niche and fitness differences (Kraft *et al*., 2015; Letten *et al*., 2017; Pérez-Ramos *et al*., 2019; Gallego *et al*., 2019; Spaak & De Laender, 2021). Resource competition, apparent competition and competition for mutualists act similarly on species coexistence (Chesson, 1990; Chesson & Kuang, 2008; Kandlikar *et al*., 2019). Coexistence appears to be limited by increasing fitness differences as species richness increases (Spaak *et al*., 2021b).

However, modern coexistence theory predominantly focuses on species from a single trophic level competing either for abiotic resources or interact phenomenologically (Narwani *et al*., 2013; Godoy & Levine, 2014; Germain *et al*., 2016). While some work on competition for biotic resources exists (Chesson, 1990; Letten *et al*., 2017), these biotic resources never have interactions with other resources or compete among themselves for resources at a lower trophic level. As a result, we have little understanding of coexistence mechanisms in the highest trophic level or, more generally, how the entire community coexists (Godoy *et al*., 2018). Additionally, modern coexistence theory is often applied to models with phenomenological or linear terms describing species interactions, excluding higher order interactions (Chesson, 2018; Germain *et al*., 2016; Pérez-Ramos *et al*., 2019) but see (Spaak *et al*., 2021a; Shoemaker *et al*., 2019; Spaak & De Laender, 2021; Spaak *et al*., 2021b; Singh & Baruah, 2020; Letten & Stouffer, 2019). Consequentially, we have only limited understanding of the drivers and mechanisms of species coexistence in communities including multiple trophic levels and non-linear species interactions.

In this paper, we investigate the mechanisms and trait differences that allow coexistence of species in an empirically parameterized model for an aquatic foodweb consisting of nutrients, phytoplankton and zooplankton. We excluded higher trophic levels because we lacked empirical data for their parametrization. We then ask whether, or to what extent, modern coexistence theory helps us to understand our results, or leads to additional insights that we would not otherwise have obtained.

We chose planktonic food webs for our model system because the mechanisms of how phytoplankton compete for resources and how zooplankton graze on phytoplankton are well understood, and have been studied and modeled for many decades (e.g., Tilman *et al*., 1982; Brun *et al*., 2016). Additionally, the models can be parameterized empirically using databases of measured phytoplankton and zooplankton traits. Using these trait databases, we can generate a species pool consisting of many thousands of hypothetical phytoplankton and zooplankton species, and then search for generalities as opposed to special cases (Litchman & Klausmeier, 2008; Litchman *et al*., 2013). Understanding planktonic foodwebs is also practically important, as they forms the basis of every aquatic foodweb and are responsible for roughly 50% of the worlds primary production (Field, 1998).

We investigated which of the mechanisms present in the community model, e.g. nutrient competition, competition for light, or predation, are most important for species richness. Additionally we wanted to understand the relative importance of different trait trade-offs for governing species richness, e.g. nutrient affinity trade-offs, growth-defense trade-offs, quality-quantity or gleaner-opportunist trade-offs. Finally, we applied methods from modern coexistence theory to gain a more general understanding of the mechanisms driving species richness. We found that species richness was primarily driven by the trophic interaction between phytoplankton and zooplankton. Competition of phytoplankton for nutrients was less important for species richness. Decreased zookplankton species richness was mostly associated with increased fitness differences. However, in many cases we could not compute the niche and fitness differences, as the zooplankton interaction is equivalent to the interaction between obligatory mutualists. Therefore, coexistence in higher trophic levels is governed by a mechanism not present in lower trophic levels and modern coexistence theory, in its current form, offers only partial insight.

## Methods

### Growth dynamics

We model an aquatic food web with two trophic levels, phytoplankton and zooplankton (Figure 1). We use superscript *P* for phytoplankton related terms and superscript *Z* for zooplankton related terms. Additionally, a subscript *i* indicates the identity of a focal phytoplankton species, *n* as a summation index of phytoplankton species and *j* for the identity of a zooplankton species. Notation for the model is defined in the text, and summarized in Table 1.

**Table 1:** Summary of model notation and allometric scalings of plankton traits (Finkel *et al*., 2010; Brun *et al*., 2016; Ehrlich *et al*., 2020).

| Variable | Description | Unit | Allometric scaling |
| --- | --- | --- | --- |
| Environment |  |  |  |
| $d$ | dilution rate | $d^{-1}$ | |
| $S_N, S_P$ | Resource supply | $\mu\text{mol } L^{-1}$ | |
| $I$ | Incoming light | $\mu\text{mol quanta } m^{-2}s^{-1}$ | |
| $z_m$ | Epilimnion depth | $m$ | |
| Phytoplankton |  |  |  |
| $V_i^P$ | Volume | $\mu m^3$ | |
| $N_i^P$ | Phytoplankton density | $cells \text{ } ml^{-1}$ | |
| $\mu_i^P$ | Maximum growth rate | $d^{-1}$ | $\sim (V_i^P)^{-0.25}$ |
| $k_{iP}^P, k_{iN}^P$ | Half-saturation for N and P | $\mu\text{mol } L^{-1}$ | $\sim (V_i^P)^{0.5}$ |
| $c_{iP}, c_{iN}$ | Resource uptake | $\mu\text{mol cell}^{-1}d^{-1}$ | $\sim (V_i^P)^{0.667}$ |
| $a_i$ | Absorption coefficient | $mm^2 \text{ cell}^{-1}$ | $\sim (V_i^P)^{0.77}$ |
| $k_{iL}^P$ | Half-saturation for light | $\mu\text{mol quanta } m^{-2}s^{-1}$ | $\sim (V_i^P)^{-0.08}$ |
| $w_i^P$ | Nutritional value of phytoplankton | $\mu\text{mol cell}^{-1}$ | $\sim (V_i^P)^{0.80}$ |
| $e_i^P$ | Edibility of phytoplankton | 1 | $\sim (V_i^P)^{-0.21}$ |
| Zooplankton |  |  |  |
| $N_j^Z$ | Zooplankton density | $ind \text{ } L^{-1}$ | |
| $V_j^Z$ | Zooplankton size | $mg \text{ } C$ | $V_j^Z$ |
| $\mu_j^Z$ | Maximum growth rate | $d^{-1}$ | - |
| $c_j^Z$ | Clearance rate | $ml \text{ } h^{-1}ind^{-1}$ | $\sim (V_j^Z)^{1.0}$ |
| $m_j^Z$ | Mortality rate | $d^{-1}$ | $\sim (V_j^Z)^{0.092}$ |
| $k_j^Z$ | Half-saturation for Nutrients | $\mu\text{mol } R \text{ } ind^{-1} \text{ } h^{-1}$ | $\sim (V_j^Z)^{1.0}(\mu_j^Z)^{1.0}$ |
| Joint variables |  |  |  |
| $h_{ji}$ | Handling time | $h \text{ cell}^{-1}ind$ | $\sim (V_i^P)^{1.0}(V_j^Z)^{-0.61}$ |
| $s_{ji}$ | Selectivity | 1 | see text |

**Figure 1:**
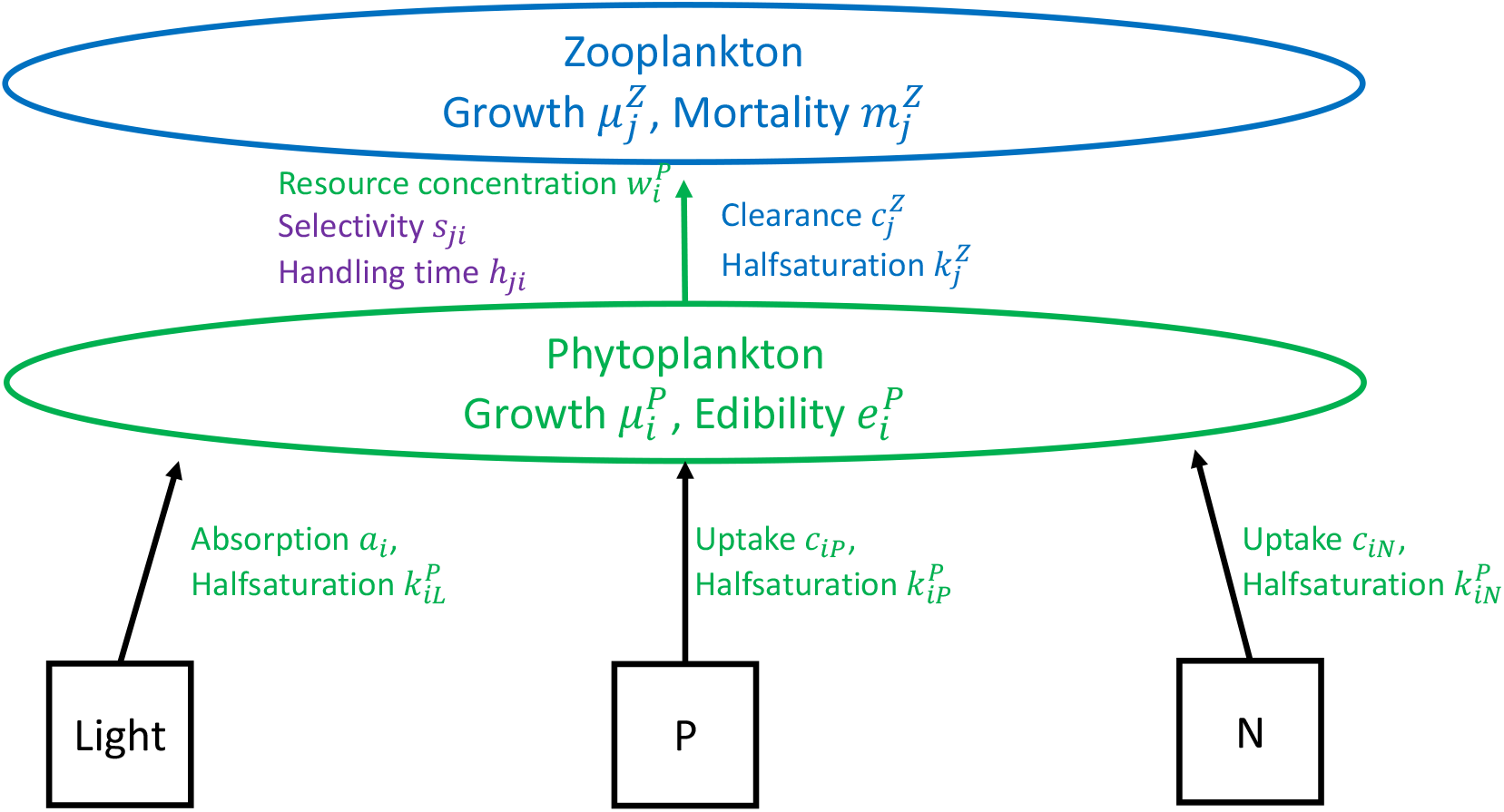
Each phytoplankton species consumes three essential resources; light, phosphorus and nitrogen. Their growth rates saturate in resource availability (Holling type 2). The growth rates of the phytoplankton are governed by resource uptake traits (*a*_*n*_, *c*_*iN*_ and *c*_*iP*_), half-saturation constants (*k*_*iL*_, *k*_*iP*_ and *k*_*iN*_) and their maximum growth rates 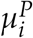. The consumption of phytoplankton is governed by their edibility 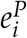, the zooplankton’s clearance rate 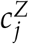, as well as the handling time *h*_*ji*_ and the selectivity *s*_*ji*_, which depend on both the identity of each phytoplankton and each zooplankton species. Given this consumption, the zooplankton growth rate is governed by the resource concentration of the phytoplankton 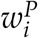, the half-saturation 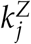 and the maximum growth rate 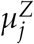 of the zooplankton. Finally, each zooplankton has a mortality rate *m*_*j*_.

We assume that phytoplankton compete for the essential resources nitrogen, phosphorus, and light (Tilman *et al*., 1982; Huisman & Weissing, 1994). The equations for these three essential resources are, respectively:

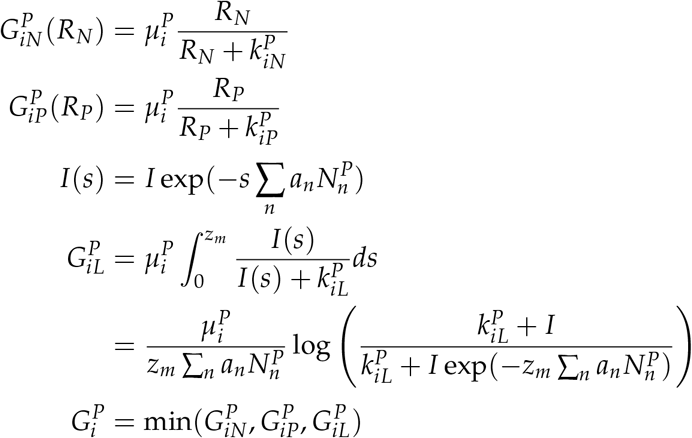

where 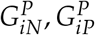 and 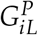 correspond to the growth rate of the phytoplankton if they were limited by nitrogen, phosphorus or light. 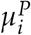 is the maximum growth rate [*day*^*−*1^], *R*_*N*_ and *R*_*P*_ are the concentration of nitrogen and phosphorus 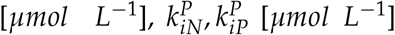 and 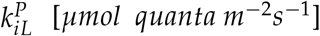 are the half saturation-constants for growth dependent on nitrogen, phosphorus and light. *s* is the depth [*m*], *z*_*m*_ is the maximum mixing depth or epilimnion depth [*m*], *a*_*n*_ is the light absorption coefficient of species *n* [*mm*^2^*cell*^*−*1^], and *I* is the incoming light intensity at surface level [*μmol quanta m*^*−*2^*s*^*−*1^]. The actual growth rate of the phytoplankton 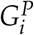 is the minimum of these three resource-specific growth rates.

The resource dynamics are given by

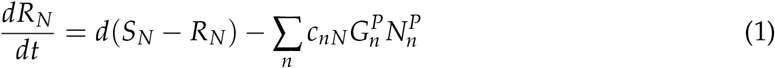

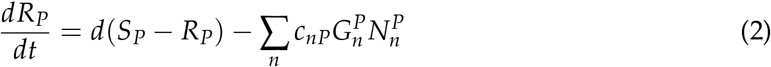

where *d* is the dilution rate of the system [*day*^*−*1^], *S*_*N*_ and *S*_*P*_ are the incoming resource concentrations of nitrogen and phosphorus [*μmol L*^*−*1^], *c*_*nP*_ and *c*_*nP*_ are the maximal resource uptake traits for nitrogen and phosphorus [*μmol cell*^*−*1^*day*^*−*1^], and 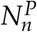 is the density of phytoplankton [*cells ml*^*−*1^]. The phytoplankton only consume the resources used directly for growth; we exclude any internal storage of resources, or, equivalently, we assume that the internal resource dynamics of the phytoplankton are fast, so that internal resource store is a function of the current nutrient uptake rate.

The zooplankton consume the phytoplankton. Each zooplankton has a clearance rate 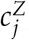, which describes the rate at which it searches for phytoplankton [*ml h*^*−*1^*ind*^*−*1^]. Zooplankton select for phytoplankton based on their size, described by a selectivity coefficient *s*_*ji*_ (dimensionless). Zooplankton *j* therefore encounter a phytoplankton *i* at rate 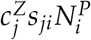, which are handled in time *h*_*ji*_ [*h cell*^*−*1^*ind*]. Additionally, certain phytoplankton are defended, making them less edible and harder to digest, represented by dimensionless edibility coefficient 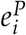 (Ehrlich *et al*., 2020). A zooplankton consumes phytoplankton at the rate (Branco *et al*., 2020)

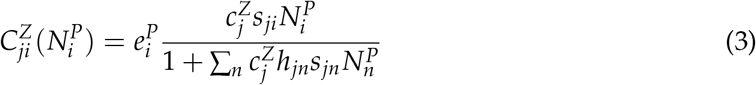

The dynamics of phytoplankton are given by

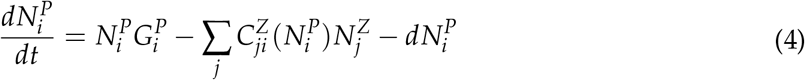

where the three terms stand for growth, grazing, and dilution of the system.

Zooplankton take up resources via consumption, so resource uptake depends on the nutritional value 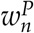 of the phytoplankton [*μmol R cell*^−1^]. 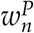 denotes the general nutritional value of the phytoplankton and we do not distinguish between phosphorus, nitrogen, or other potentially limiting resources. We assume that the nutritional value of phytoplankton does not depend on external nutrient concentrations, as the stoichiometry of the phytoplankton is approximately constant. Rather, external nutrient concentrations only affect phytoplankton growth rates. The zooplankton growth rates are given by

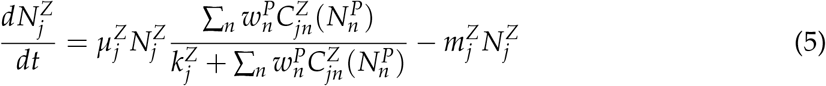

where 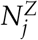 is the density of zooplankton 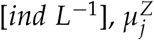 is the maximum growth rate of the zooplankton 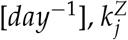 is the half-saturation constant for zooplankton growth [*μmol R ind*^*−*1^ *h*^*−*1^], and 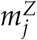 is the mortality rate of zooplankton *j* [*day*^*−*1^].

### Allometric scaling and parameter definitions

The phytoplankton traits 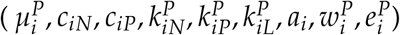 and zooplankton traits 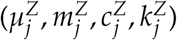 were randomly drawn from a multivariate lognormal distribution. All the multivariate lognormal distributions were fitted to empirical data from the literature (Edwards *et al*., 2012; Brun *et al*., 2016; Ehrlich *et al*., 2020). For each trait *T*, we first fitted a log-normal distribution to the empirically measured data, fitting mean *μ*_*T*_ and standard deviation *σ*_*T*_, i.e. log(*T*) *∼ 𝒩* (*μ*_*T*_, *σ*_*T*_) (Figure S9 and S10, diagonal). Unfortunately, the datasets did not contain sufficient data-points to estimate the correlations between the log-distribution of the traits.

We estimated the correlation of the lognormal trait distributions using allometric scaling. We compiled allometric scaling parameters for each of the traits (Finkel *et al*., 2010; Litchman *et al*., 2007; Brun *et al*., 2016; Ehrlich *et al*., 2020), i.e. log(*T*) *∼ β*_*T*_ log(*V*), where *V* is the volume of the phyto- or zooplankton. We then computed the correlation between two traits *T*_1_ and *T*_2_ as

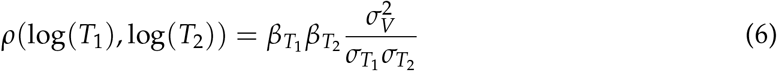

where *σ*_*V*_ is the standard deviation of log(*V*).

Given the correlation matrix *ρ* of log plankton traits, we computed the covariance matrix Σ = *σρσ*^*T*^, where *σ* is the vector of standard deviations of the log trait distributions. The log plankton traits were assumed to have a multivariate normal distribution with mean *μ*_*T*_ and covariance matrix Σ. This method allowed us to define all species-specific traits 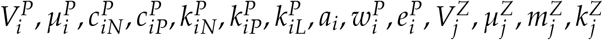 and 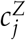 (Figure S9 and S10, off-diagonal). For the half-saturation constant 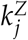 we found that 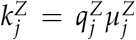, where 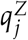 is the minimal resource concentration of zooplankton *j* [*μmol R mg C*^*−*^1] (see appendix S2).

The handling time *h*_*ji*_ and the selectivity *s*_*ji*_ depended on the traits of both species in a phytoplankton-zooplankton pair and therefore could not be determined with the above method. The handling time decreases with zooplankton size and increases with phytoplankton size and was defined as 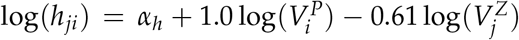 (Branco *et al*., 2020; Uye, 1982). Selectivity *s*_*ji*_ was a decreasing function of the difference between phytoplankton size and preference of size by zooplankton. We set the size preference of the zooplankton to 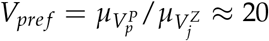, i.e. a zooplankton of mean size would prefer phytoplankton of mean size (Berggreen *et al*., 1988). Given this, we defined the relative selectivity of zooplankton to be

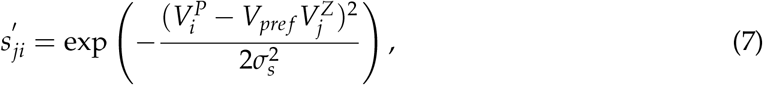

where *σ*_*s*_ is the selectivity breadth, which was set to 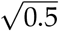 (Branco *et al*., 2020). We then normalized these relative selectivities such that their total for each zooplankton is 1, i.e. 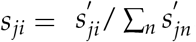

### Simulations

To assess the relationship between trait distributions and community species richness, we conducted simulations that mimic community assembly. We first generated 1000 species pools each consisting of 20 phytoplankton and 20 zooplankton species. For each species pool, we started community assembly with one randomly chosen phytoplankton species at its monoculture equilibrium density and one randomly chosen zooplankton species at low density. We then simulated the community dynamics for one year, removed any species that went extinct (below 0.01% of total community biomass), and introduced one new phytoplankton and zooplankton species at low density. Our analyses are based on the resulting species richness after 20 years (longer times did not increase species richness, Appendix S1, Fig. S1). Strictly speaking, we assessed co-occurrence rather than coexistence, as we simply observed the presence of species at the end of the simulation. However, often the sub-communities with one species removed did not form a stable equilibrium, so that computing invasion growth rates was not possible (see *Results, Niche and Fitness Differences*, below).

We assessed the importance of each trait for species richness by simulating community assemblies of plankton with altered traits. Specifically, we independently altered the mean *μ*_*T*_ or the standard deviation *σ*_*T*_ of the log distribution of each trait distribution. We increased or decreased *μ*_*T*_ by one standard deviation *σ*_*T*_, and we increased or decreased the standard deviation *σ*_*T*_ by a factor of 4 (corresponding to a change in variance by a factor of 2). With these altered traits we performed the same community assembly process (Fig.S2).

We investigated the effect of the standard deviation of the handling time *h*_*jn*_, by setting 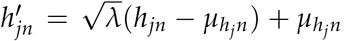, where *λ* is the factor by which we increased or decreased the standard deviation and 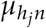 is the mean of the handling time. We investigated the effect of standard deviation of the selectivity by setting 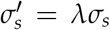. Because the total selectivity of a zooplankton must sum to 1 (Σ_*n*_ *s*_*jn*_ = 1) we did not investigate the effect of mean selectivity on species richness.

We investigated the effect of trait trade-offs by altering the correlation *ρ*(*T*_1_, *T*_1_) between two traits. The correlation between two traits cannot be set to any arbitrary value in [-1,1], rather the resulting correlation matrix *ρ* must be semi-positive definite. We simulated communities with the minimal and maximal possible value for correlation, which depends on the specific traits involved.

### Niche and fitness differences

To better understand how the traits affected coexistence, we computed niche and fitness differences for zooplankton species competing for phytoplankton using the method of Spaak *et al*. (2021c). To compute niche and fitness differences we computed the intrinsic, the invasion and the no-niche growth rates. These were defined as the growth rates of the zooplankton “invading” three different scenarios. The intrinsic growth rate, denoted *μ*_*j*_ describes the growth rate of a zooplankton invading an empty community, where no zooplankton are present and phytoplankton are at their equilibrium. Note that the intrinsic growth rate *μ*_*j*_ is lower than the maximal growth rate 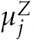, as 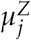 is the growth rate when resource abundance for the zooplankton is infinite. The invasion growth rate *r*_*j*_ is the growth rate of a zooplankton invading a community with the competitor zooplankton at their equilibrium densities and phytoplankton at their corresponding densities. Finally, the no-niche growth rate *η*_*j*_ is the growth rate of the zooplankton invading a community with itself as a resident, but at a density equivalent to the combined equilibrium density of its competitors.

We only computed niche and fitness differences for the zooplankton species, whereas phytoplankton were considered only as resources. We could not compute the niche and fitness differences for competing phytoplankton, as this would have required a function for the growth rates of the phytoplankton 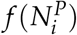 which only depends on the densities of the phytoplankton 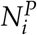 (Spaak & De Laender, 2020). However, this function is not defined, as for a given phytoplankton density 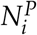, the zooplankton growth rates cannot be set to zero, and hence their density is not defined.

Given these growth rates, we defined niche and fitness differences as

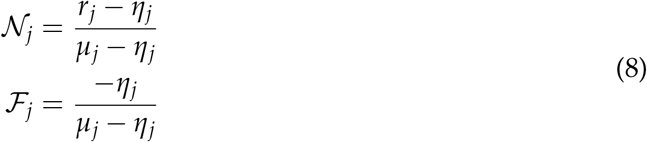

Importantly, these definitions of niche and fitness differences depend on the invasion growth rate and can therefore only be computed for communities in which both species in monoculture reach a stable, nonzero equilibrium density. However, this was only the case for a small fraction of the investigated communities (see Results).

## Results

In simulations with unaltered trait distributions (i.e., the distributions estimated from empirical measurements), phytoplankton species richness ranged from 1 to 5 with an average of 2.4, and zooplankton species richness ranged from 0 to 5 with an average of 2.2 (Fig. 2). Phytoplankton and zooplankton richness were strongly correlated (*ρ* = 0.71); roughly 50% of communities had equal phytoplankton and zooplankton species richness, and zooplankton richness exceeded phytoplankton richness in only 15% of the communities.

**Figure 2:**
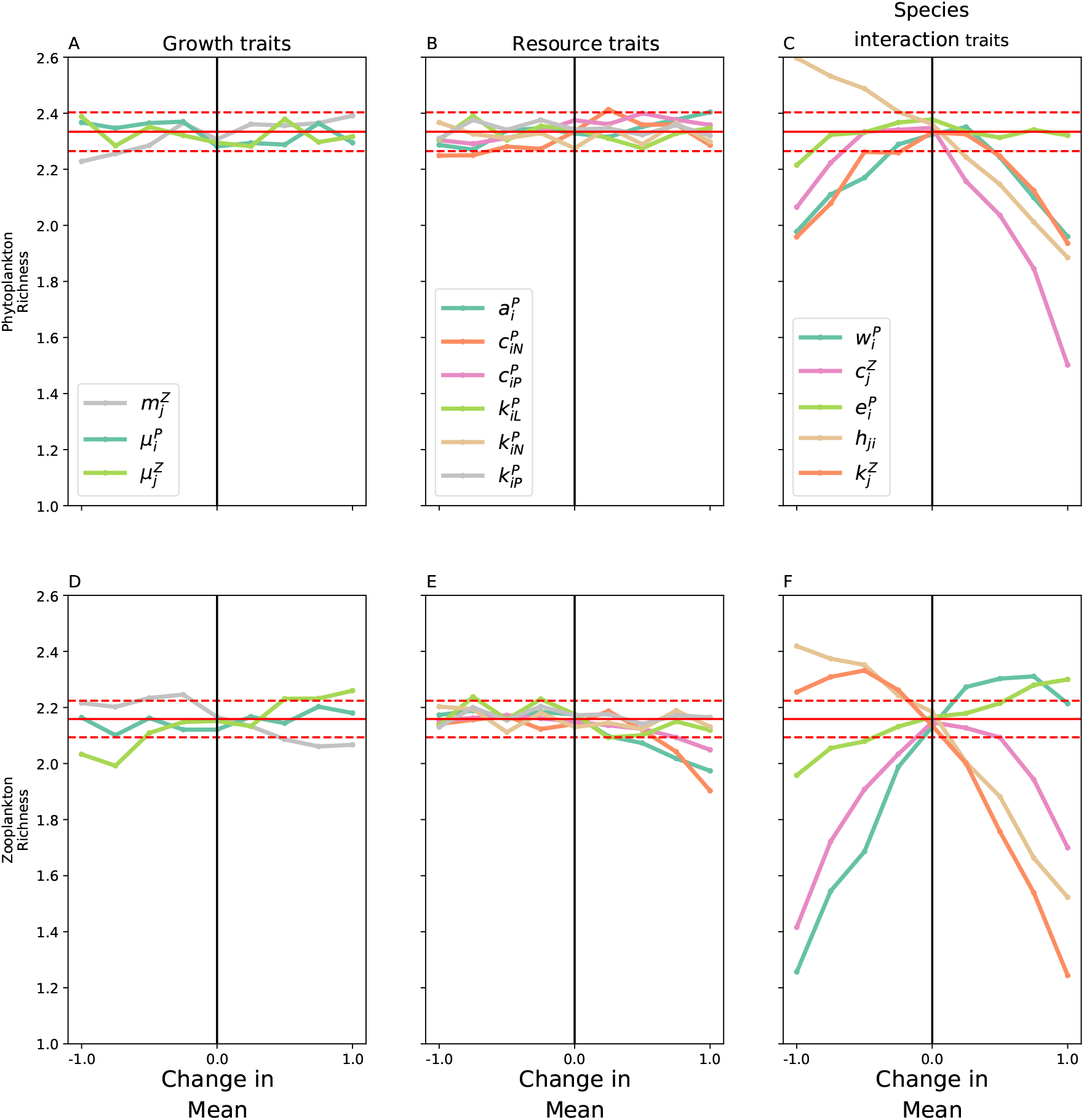
We changed the means of growth traits (A,D), resource traits (B, E) and species interaction traits (C,F) and investigated their effect on phytoplankton (A,B,C) and zooplankton (D,E,F) species richness. A,D: The mean values of the growth traits (A,D) had no effect on species richness. B,E: Similarly, the mean values of the traits governing the competition of phytoplankton for resources had little effect on species richness. C,F: Conversely, altering the traits governing the interaction between phytoplankton and zooplankton had a strong effect on species richness. Many of these trait changes decreased the viability of the zooplankton and led to zooplankton starvation. Decreased zooplankton richness decreases the stabilizing effect of zooplankton on phytoplankton which then decreases phytoplankton richness. A-F: Black vertical line indicates the empirical values of the traits. Red horizontal line show the mean and 99% confidence interval of the respective values for communities generated with the empirical trait values, to indicate the sampling variability of species richness estimates.

Based on resource competition theory, zooplankton species richness should not exceed the number of phytoplankton species, as the number of coexisting species cannot exceed the number of limiting factors at a stable equilibrium. However, the system does not always reach a stable equilibrium, rather the phytoplankton and zooplankton induce predator-prey cycles which could lead to coexistence via storage effects or relative non-linearities (Armstrong & McGehee, 1976, 1980; Litchman & Klausmeier, 2001; Huisman & Weissing, 1999). Moreover, we did not assess whether the zooplankton actually coexisted or whether the time to competitive exclusion was very long (see below).

We did not observe strong selection for most traits (except 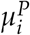), as shown by the close match between the trait distribution of surviving species and the initial trait distributions (Fig. S8). Similarly, selection did not have a strong effect on the correlation matrix of the trait distribution (Fig. S8).

Simulations with altered trait distribution showed that changes in the mean of many trait distributions had little or no effect on species richness (Fig. 2, A,B, D and E). Traits associated with species growth rates (*μ*^*P*^, *μ*^*Z*^ and *m*_*Z*_) were unimportant for species richness (A,D). Similarly, phytoplankton resource uptake traits (*c*_*n*_, *c*_*p*_ and *a*) as well as half-saturation constants for these resources (*k*_*n*_, *k*_*p*_ and *k*_*l*_) had only minor effects on species richness. When these traits had very large values, species richness of zooplankton declined (E), likely because many zooplankton species did not have enough phytoplankton to consume, as high resource uptake as well as high half-saturation constant implies lower equilibrium densities of phytoplankton.

Changes in traits regulating the phytoplankton-zooplankton interactions had the strongest effect on species richness (Fig. 2, C, F). This matches findings from observational data in a temperate lake (Merkli, 2021). The effects of these trait changes on species richness are likely due to a combination of the following three explanations:

First, the trait changes affected the amount of resources taken up by zooplankton. If this amount was to low, e.g. because consuming one phytoplankton cell takes too long (high handling time *h*_*ji*_, brown line, Fig. 2 C and F), then zooplankton starved and species richness droped. Conversely, if this amount increased, then more zooplankton had sufficient food to survive. This is a possible explanation for the effects of handling time *h*_*ji*_ and edibility 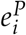 on species richness. However, this explanation ignores competition between zooplankton and leads to the prediction that the trait changes monotonically affect species richness.

Second, the trait changes affected the underlying trait trade-offs and allowed the creation of super species. This may explain why changes in three of the traits showed a unimodal effect on species richness (nutritional value 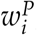, half saturation constant 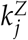 and clearance rate 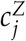). For example, the positive correlation between half saturation 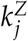 and clearance rate 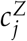 created a gleaner-opportunist trade-off. However, if all species had increased clearance rates, then growth was not limited by the amount of phytoplankton consumed (which was driven by the clearance rate), but only by the half-saturation constant, which effectively destroyed the trade-off.

Third, phytoplankton species richness follows the species richness of zooplankton, because phytoplankton richness is maintained by the trophic interaction with zooplankton. Consequentially, a decline of zooplankton species richness decreased the stabilizing effect of zooplankton on phytoplankton, and their richness declines as well (Fig. S6). This explains why resource competition traits had little effects on phytoplankton richness, as well as the strong correlation between phytoplankton and zooplankton species richness. It also explains why effects of trait changes more strongly affected zooplankton richness than phytoplankton richness.

Changing trait correlations (Fig. 3) confirmed our conclusions from the effects of changes in trait means. First, many trait correlations had no strong effect on species richness (Fig. S5). Second, many of the trait correlations which had a strong effect on richness were linked to the trophic interactions between phytoplankton and zooplankton. Third, zooplankton species richness was more sensitive to changes in trait correlations.

**Figure 3:**
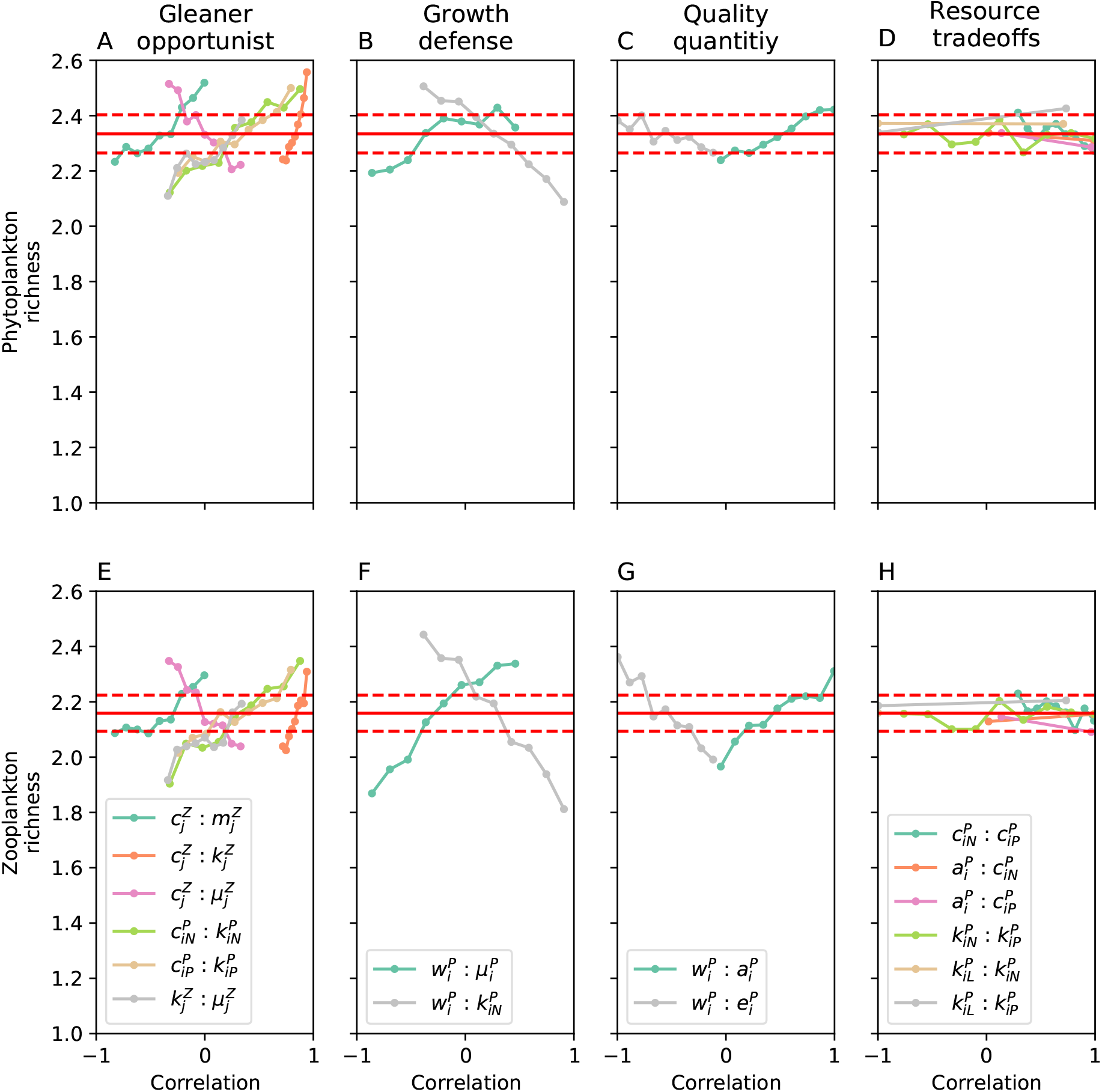
We changed the trade-offs (correlation matrix) between traits and investigated the effects on phytoplankton (A-D) and zooplankton (E-H) species richness. The trade-offs with the strongest effects are grouped into gleaner-opportunist (A,E), growth-defense (B,F) and super-resource (C,G) trade-offs. Interestingly, none of the resource competition trade-offs had a strong effect on species richness (D,H). Generally, zooplankton richness (E-H) was more sensitive than phytoplankton richness (A-D) to changes in trade-offs, similar to the changes in mean (Fig. 2). A-H: Red horizontal line show the mean and 99% confidence interval of the respective values for communities generated with the empirical trait values. The correlation cannot be chosen freely for each trade-off, rather the maximal and minimal possible values for each trait pair are determined by the requirement that the entire trait correlation matrix must be positive definite (see Methods).

Six of the ten trade-offs with the strongest effect can be categorized as gleaner-opportunist trade-offs, i.e. one species benefits from high resource availabilities and the other from low resource availabilities (Fig. 3 A,E).

Two of these trade-offs are competition for nitrogen (green line) and phosphorus (brown line), the other four concerned the trophic interactions. The other trade-offs all concerned the nutritional value 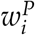 of the phytoplankton. Two of these are conceptually similar to a growth-defense trade-off (B,F). Low nutritional value 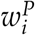 does not provide immediate protection against predation, however it limits the growth rates of the predator and therefore protects against future predation. High intrinsic growth rate 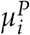 or low half saturation constant 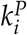 both contribute to high growth rates, so these trade-offs are therefore similar to the familiar growth-defense trade-offs. However, the growth defense trade-off between edibility 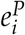 and intrinsic growth rate 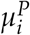 did not have a strong effect on species richness.

The last two are trade-offs between high quality food (i.e. high nutritional value 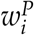), and high quantity food, either high abundance because of low absorption rates 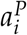 or high edibility 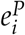. Reducing the strength of this trade-off will create phytoplankton having both high quality and quantity. Zooplankton species that consume these abundant high quality phytoplankton will have a competitive advantage over zooplankton consuming low quality and quantity phytoplankton, and that competitive advantage will decrease zooplankton richness.

Interestingly, none of the six possible resource trade-offs had a strong effect on species richness (Fig. 3 D,H). This supports our hypothesis that differences among phytoplankton species in resource requirements, or response to limiting resources, are not important drivers of species richness in this community.

### Niche and fitness differences

Changes in fitness differences were the underlying cause for the change in zooplankton species richness for most changes in trait distributions (Fig. 4 D). The trait changes that lead to starving zooplankton (increasing half saturation constant *k*_*Z*_, increasing handling time *h*_*j*_ *p*, decreasing resource availability *R*_*P*_ and decreasing clearance rate *c*_*Z*_) all increased fitness differences between the competing zooplankton (Fig. S3 and S4). The closer a zooplankton was to the starvation boundary, the lower its fitness and the stronger fitness differences became, explaining the decrease in species richness. Similarly, relaxing gleaner-opportunist trade-offs created super species with increased fitness (Fig. 3 E); the growth-defense trade-offs and the quality-quantity trade-offs create super resources, and zooplankton consuming these super resources had a fitness advantage (Fig. 3 F,G). Niche differences between zooplankton were not correlated with zooplankton species richness (Fig. 4 C). Similarly, niche and fitness differences between competing zooplankton were not good predictors of phytoplankton species richness.

**Figure 4:**
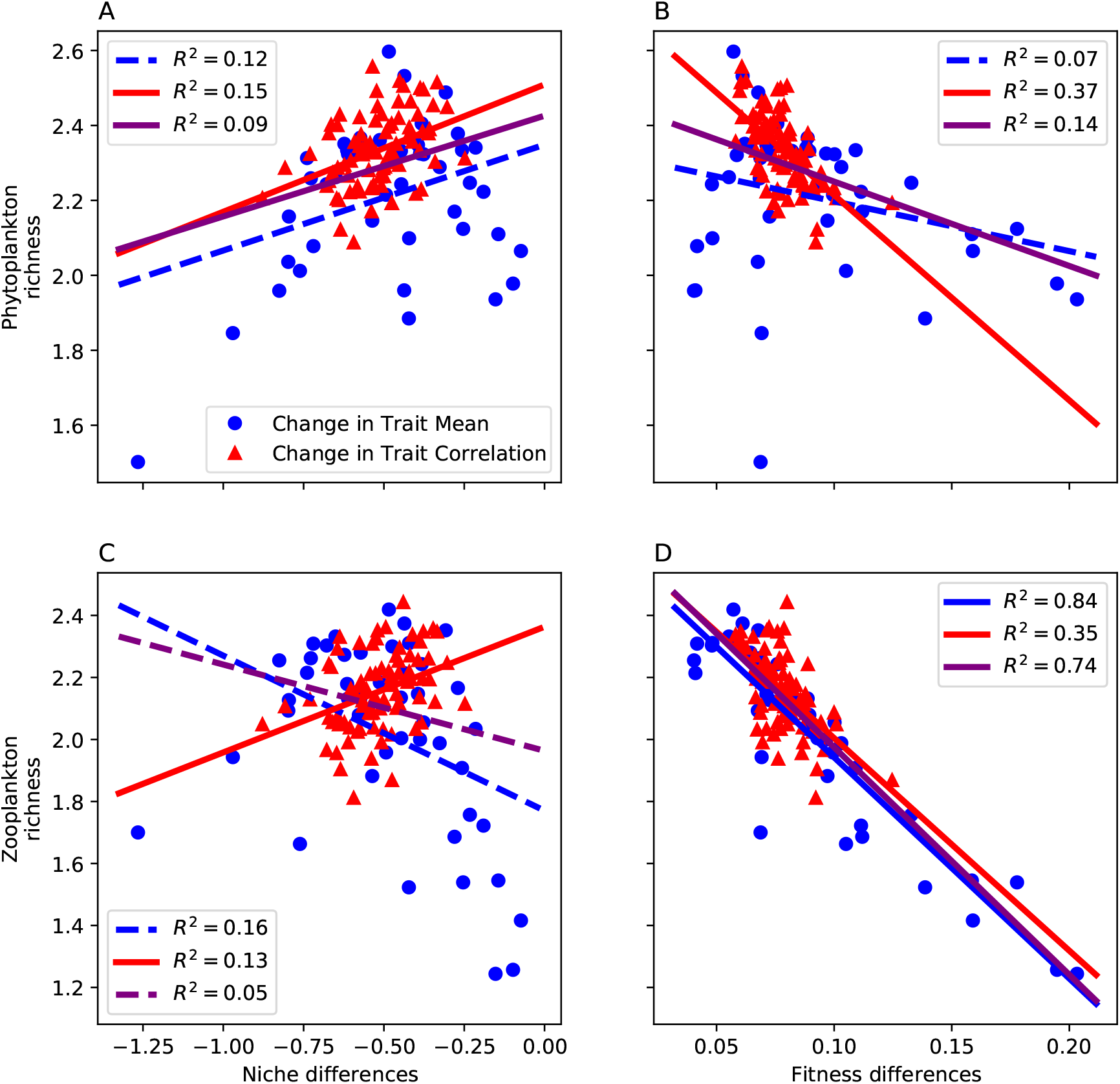
Overall, niche differences were not strongly correlated with phytoplankton (A) or zooplankton richness (C). D: Fitness differences are strongly correlated with zooplankton richness. B: The correlation between fitness differences and phytoplankton richness likely stems from the correlation between phytoplankton and zooplankton richness, and not from a direct cause of fitness differences on phytoplankton richness. Blue dots show average species richness from changes in mean traits, i.e. corresponding to Figure 2 C or F versus the niche or fitness differences from changes in mean traits, (Fig. 2 H or I). Red triangles show the corresponding data for changes in trade-offs (Fig. 3). Lines show the best linear fit based on least *R*^2^, purple line show the best linear fit for all data. Dashed lines indicate that linear fit was not significant at the *p* = 0.01 level.

However, the relevance of these findings is not entirely clear because we were only able to compute niche and fitness differences for a small fraction of all assembled communities (*∼*10%). In roughly 40% of communities, one of the two zooplankton had no stable equilibrium density in monoculture, but rather an equilibrium density distribution. Computation of niche and fitness differences is technically still possible for these communities, but turned out to be computationally very expensive and imprecise. To compute niche and fitness differences, one first has to compute the equilibrium density distribution of both species, which often required simulations of more than 5000 days per species, as the densities over time were often very chaotic. Given this distribution, one has to numerically solve the equation 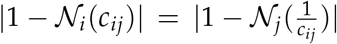 for *c*_*ij*_, however, for each iteration of the numerical solver one has to recompute the equilibrium distribution given the new *c*_*ij*_, which was computationally very time-consuming. We therefore did not compute niche and fitness differences for these communities.

Additionally, in some communities one of the two zooplankton was not able to persist at all in the absence of the other zooplankton. This occurred when the two zooplankton could coexist, because of the zooplankton behaving similarly to obligatory mutualists. As an illustrative example, consider a large and a small zooplankton that predominantly feed on a large or a small phytoplankton species, respectively (Fig. 5 B). In the absence of any zooplankton, suppose that the smaller phytoplankton has an advantage in competition for the abiotic resources, and competitively excludes the larger phytoplankton species. If the larger zooplankton is an inefficient predator of the small phytoplankton and therefore cannot invade a community where only small phytoplankton are present, it will not have a monoculture equilibrium density. Consequently, we cannot compute an invasion growth rate of the smaller zooplankton invading the larger zooplankton in monoculture, and computation of niche and fitness differences is not possible.

**Figure 5:**
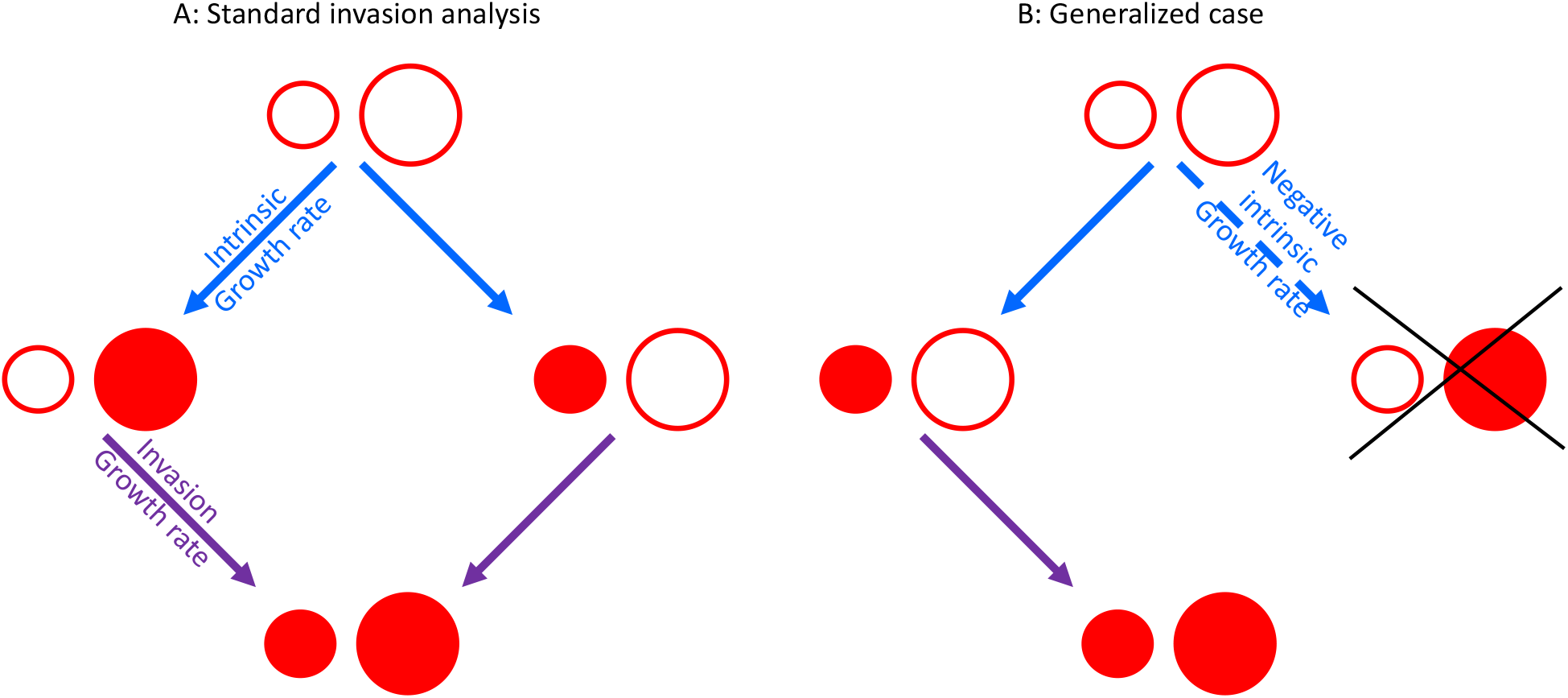
A: The standard community assembly process of a small and a large zooplankton competing for phytoplankton. Initially, both zooplankton species are absent (top) and both have a positive intrinsic growth rate. Either of the two zooplankton will arrive and successfully invade the empty ecosystem as both have a positive intrinsic growth rate (blue arrow) leading to an ecosystem in monoculture (middle). Finally, the other zooplankton species can invade the monoculture of its competitor, as both have a positive invasion growth rate (purple arrow) leading to coexistence (bottom). B: However, if we assume that the lower trophic level of phytoplankton cannot coexist in the absence of zooplankton only one of the zooplankton (here the smaller) will have a positive intrinsic growth rate, as the other zooplankton will (potentially) not have enough food. Consequentially, the larger zooplankton will not be able to survive in monoculture and not have a monoculture equilibrium density, therefore the invasion growth rate of the smaller zooplankton is not defined in the traditional sense. We can therefore not apply modern coexistence theory in its current form to this community. However, coexistence might still be possible, as the large zooplankton can invade the monoculture of the small zooplankton. This is possible because the small zooplankton reduces the competitive ability of the small phytoplankton such that the small and the large phytoplankton can coexist in presence of zooplankton.

Nonetheless, the two zooplankton might be able to coexist. The smaller zooplankton will predominantly consume the smaller phytoplankton. If we first introduce the smaller zooplankton it will decrease the density of the smaller phytoplankton and reduce its competitive strength. The larger phytoplankton will invade and persist in the presence of the smaller zooplankton. Then, the larger zooplankton will have a prey to consume and may be able to invade. Even though all species interactions result from resource competition, this scenario resembles the cases of obligatory mutualism or keystone species and cannot be analyzed using any of the existing definitions of niche and fitness differences (Spaak & De Laender, 2020; Spaak *et al*., 2021c).

## Discussion

We investigated the drivers of species richness in phytoplankton-zooplankton communities based on mechanistic species interactions and empirical trait data. We altered several parameters of the trait distributions governing phytoplankton and zooplankton growth and found that changes in traits associated with the trophic interaction between phytoplankton and zooplankton had strong effects on species richness (Fig. 2 C,F and 3A-C and E-F). However, changes in the resource competition traits or traits governing maximal growth rates had much weaker effects on species richness. We conclude that species richness in our model is primarily determined by the trophic interaction between phytoplankton and zooplankton. Our second important finding is that the changes in species richness caused by changes in trait means or correlations were largely driven by altered fitness differences (Fig. 4).

Trade-offs in resource affinities did not have a strong effect on species richness in our model (Fig. 3 D), for two reasons. First, while the potential benefits of such trade-offs on species richness are clear (Tilman *et al*., 1982; Letten *et al*., 2017; Huisman & Weissing, 1994), those benefits may be quite weak. In our model, two randomly chosen phytoplankton are unlikely to have a sufficiently strong trade-off in their resource affinities to stabilize coexist. Often, the smaller phytoplankton will out-compete its larger competitor based on resource competition alone (**?**Edwards *et al*., 2012). Second, the relative importance of a certain mechanism may depend on the presence of other mechanisms (Shoemaker *et al*., 2020a; Zepeda & Martorell, 2019; Letten *et al*., 2018). Most work on resource competition has been done in the absence of predators. When we remove predators, we do find that differences in resource uptake traits or growth rates of phytoplankton species have a small positive effect on species richness (Appendix S1, Fig. S7). However, in the presence of predators, the positive effect of resource competition on species richness is unimportant. These results demonstrate that instead of asking whether a mechanism operating in isolation *can* affect species richness, we should investigate both the magnitude and the relative importance of different mechanisms operating simultaneously. Consideration of multiple mechanisms also makes it possible to study interactions among them. In our model, resource partitioning and predation were not additive.

We found that fitness differences were correlated with zooplankton richness (Fig. 4 D) and to a lesser extent with phytoplankton richness (Fig. 4 B). Conversely, niche differences were only weakly correlated with phytoplankton or zooplankton richness. This differs from earlier findings in phytoplankton or plant communities, which mostly found that niche differences were better predictors of species richness (Narwani *et al*., 2017; Buche *et al*., 2021; Levine & HilleRisLambers, 2009; Adler *et al*., 2010). This difference may stem from many different sources, such as model complexity, the two trophic levels in our model, or our inability to compute niche and fitness differences for all communities (see next section).

Our model is based on mechanistic species interactions and empirically measured traits, yet it lacks certain key features from the natural world. First and foremost, we did not include any external fluctuations, either random or deterministic (e.g., seasonality), and as such we excluded many potential coexistence mechanisms (Chesson, 1994; Ellner *et al*., 2019; Letten *et al*., 2018; Shoemaker *et al*., 2020b; Barabás *et al*., 2018). Fluctuations would likely increase species richness and potentially affect the importance of resource competition. Second, species richness was generally low compared to natural communities. The relative importance of different coexistence mechanisms might depend on species richness. For example, Spaak *et al*. (2021a) have shown that increasing species richness increases the importance of fitness differences compared to niche differences. Third, the trophic structure was simple and excluded any higher trophic level and any mixotrophs. As we found that trophic interactions are the most relevant factor for species richness, it would be interesting to see whether higher trophic levels are even more important.

### Implications for modern coexistence theory

We were not able to compute niche and fitness differences for many of our simulated communities. This is not solely a limitation of the niche and fitness differences method we chose to apply, but more generally a limitation of modern coexistence theory in its current formulation based on invasion growth rates. Similar issues have been known for other models, such as models including Allee effects (Barabás *et al*., 2018). However, those models were specifically constructed to showcase the limitations of modern coexistence theory. In our model these difficulties are a natural consequence of the increased complexity of the underlying community. It is not solely a limitation of the model investigated here, rather similar issues are likely to emerge in any multi-trophic community, whether the model is mechanistic or phenomenological. For example, Huisman & Olff (1998) and Arsenault & Owen-Smith (2002) found that small herbivores can only persist in the presence of larger herbivores. Another typical example are keystone species, where the presence of these keystone species has large effects on the biodiversity of the ecosystem Creed (2000).

Much of the work of modern coexistence theory is based on invasion growth rates and the assumption that these invasion growth rates are indeed well-defined and give insight into coexistence (Spaak & De Laender, 2020; Barabás *et al*., 2018; Ellner *et al*., 2019; Pande *et al*., 2019; Schreiber, 2000). However, we observed communities in which the invasion growth rate is not well defined or does not exist (Figure 5). These communities behaved similarly to communities driven by Allee effects or communities driven by obligatory mutualism. It was not previously recognized that these communities can arise from simple resource competition, a well understood mechanism that is key for modern coexistence theory (Chesson & Kuang, 2008; Chesson, 1990; Letten *et al*., 2017; Letten & Stouffer, 2019; Spaak *et al*., 2021a,b).

Schreiber (2000) proposed a generalized version of invasion analysis. The traditional invasion growth rates ask whether each species can invade the sub-community from which it is absent (Figure 5 A). In contrast, the generalized invasion analysis investigates for each possible sub-community which species can invade (Figure 5 B). However, the available methods to compute niche and fitness differences (Spaak & De Laender, 2020; Spaak *et al*., 2021c; Carroll *et al*., 2011) do not yield additional insight, as they would always result in *𝒩* = 1 and *ℱ* = *±*∞. Similarly, the decomposition of the invasion growth rates by Ellner *et al*. (2019) or Chesson (2003) depend on the resident having zero growth rates (in order to allow a resident-invader comparison), however the resident species would have a negative growth rate. We therefore must expand modern coexistence theory to apply to such communities.

## Acknowledgments

We thank Francesco Pomati for comments on earlier versions of this script. J.W.S was supported by the Swiss national science foundation SNSF under the project P2SKP3 194960. S.P.E. was supported by US NSF grant DEB-1933497. P.B.A. was supported by US NSF grant DEB-1933561 (PBA).

## Appendix

### S1 Additional analyses

**Figure S1:**
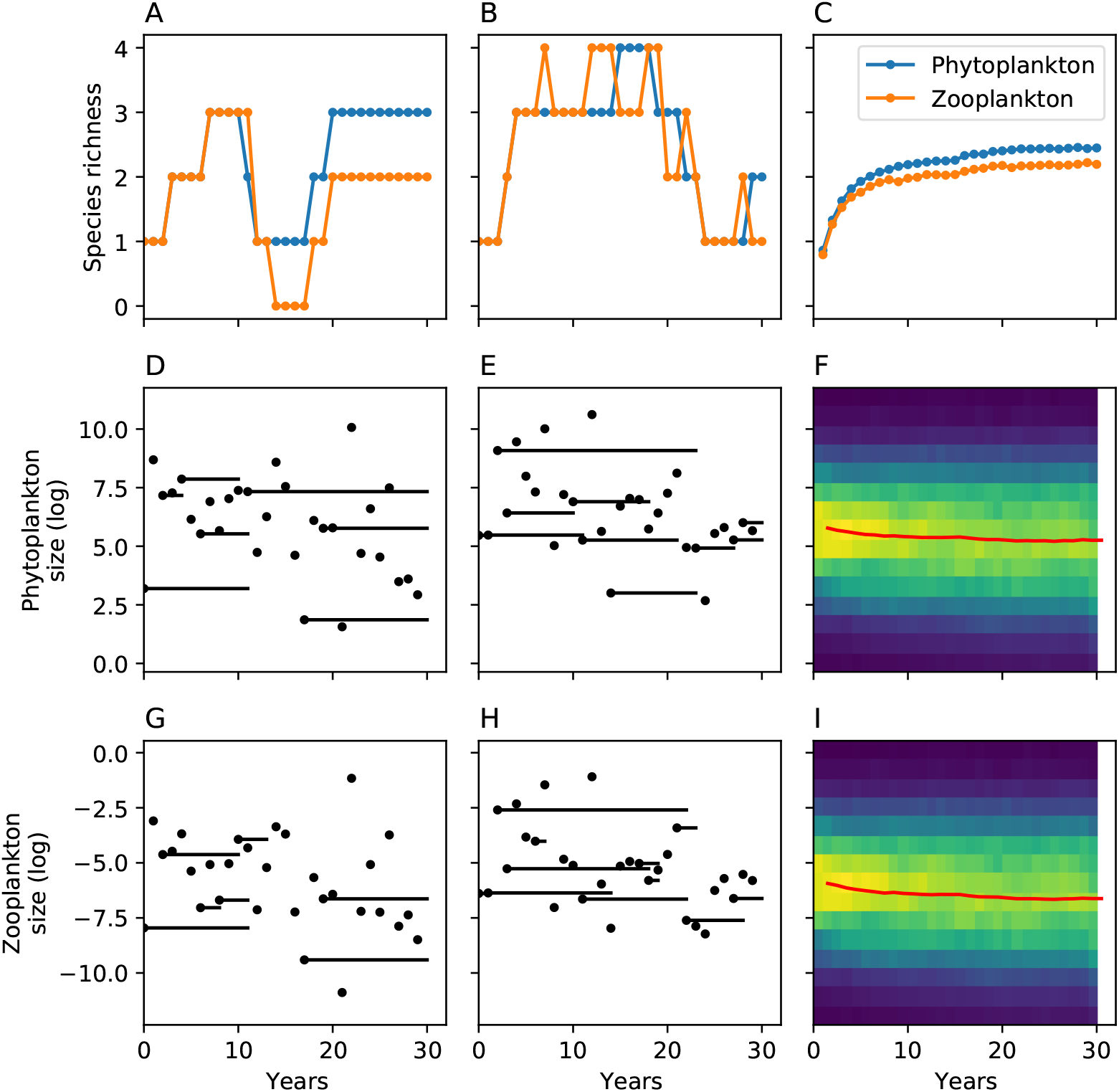
Our simulated community assembly process, for two sample communities (left and center columns) and the average over 1000 communities (right column). A,B: Species richness of phytoplankton (blue) and zooplankton (orange) over time. Richness is measured at the end of each year, and all species with density below a threshold are assumed to be extinct. C: Average species richness of phytoplankton and zooplankton species over time. Species richness is a saturating function of time, in the main text we stopped assembly after 20 years, increasing the community assembly time to 30 years would have no strong effect on species richness. D,E: Presence of phytoplankton species over time. Each dot represents when a phytoplankton species was introduced, and black lines indicate when the species remains present. y-axis indicates the size of the introduced phytoplankton species, which follows a gaussian distribution. G,H: Similar to D,E but for zooplankton species. F: For each year we plot the distribution of sizes of the present phytoplankton species. The distribution in the first year is identical to the distribution of the species pool. Average species size (red line) decreases over time slightly, as smaller species appear to have a competitive advantage. For a better comparison of the initial and final size distributions see Figure S8 E and F. I: Similar to panel F but for zooplankton species.

**Figure S2:**
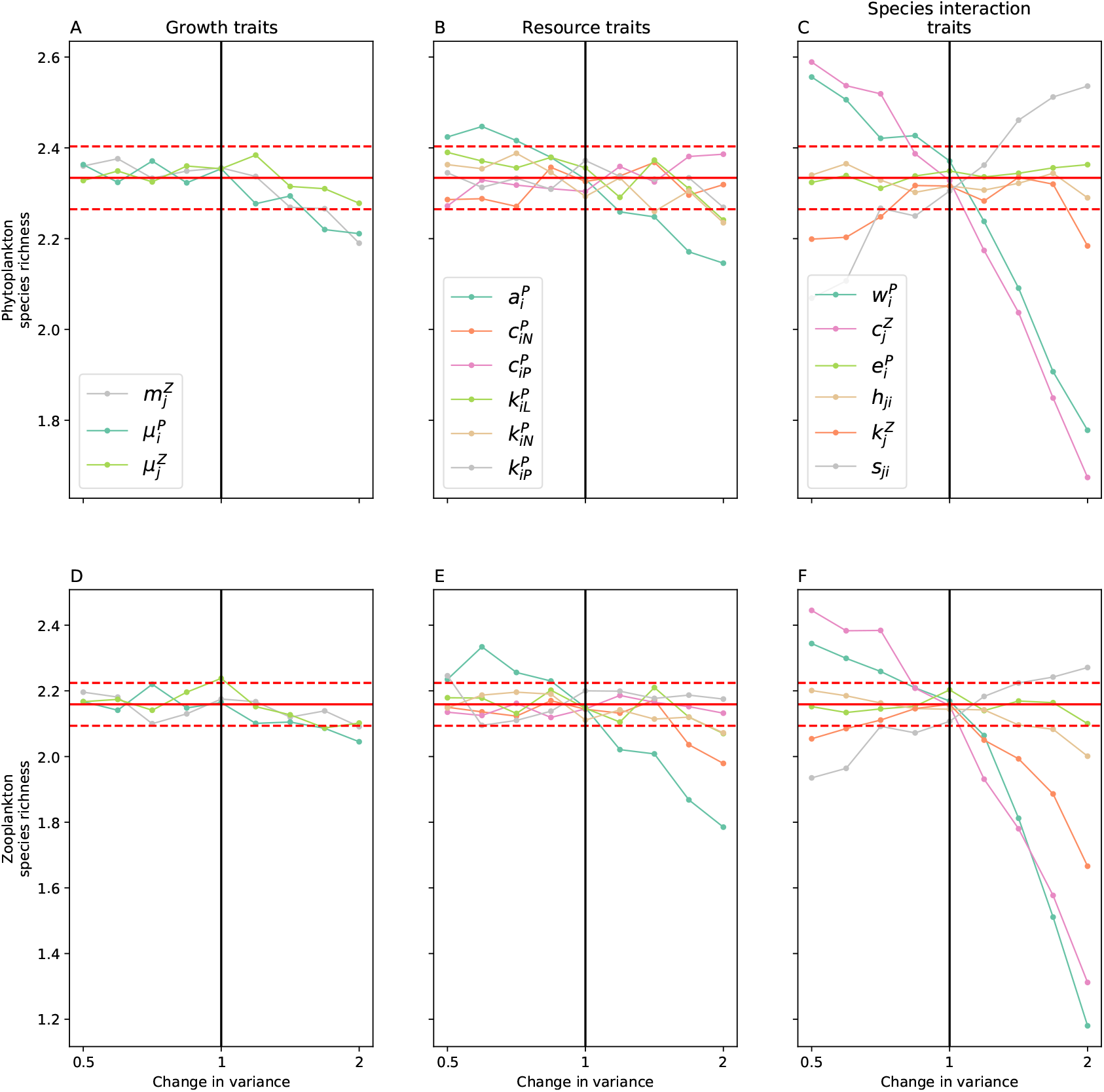
Similar to figure 2 but for changes in trait-variances, overall the results from changes in variances match the results from changes in trait means. We changed the variance of the growth traits (A,D), resource traits (B, E) and the species interaction traits (C,F) and investigated their effect on phytoplankton (A,B,C) and zooplankton (D,E,F) richness. A,D: The specific values of the growth traits (A,D) had no effect on species richness. B,E: Similarly, the specific values of the traits governing the competition of phytoplankton for resources had little effect on species richness. C,F: Conversely, altering the traits governing the trophic interaction between phytoplankton and zooplankton has a strong effect on species richness. Overall, the effect of changing the variance on the species richness are comparable to changing the mean of the traits.

**Figure S3:**
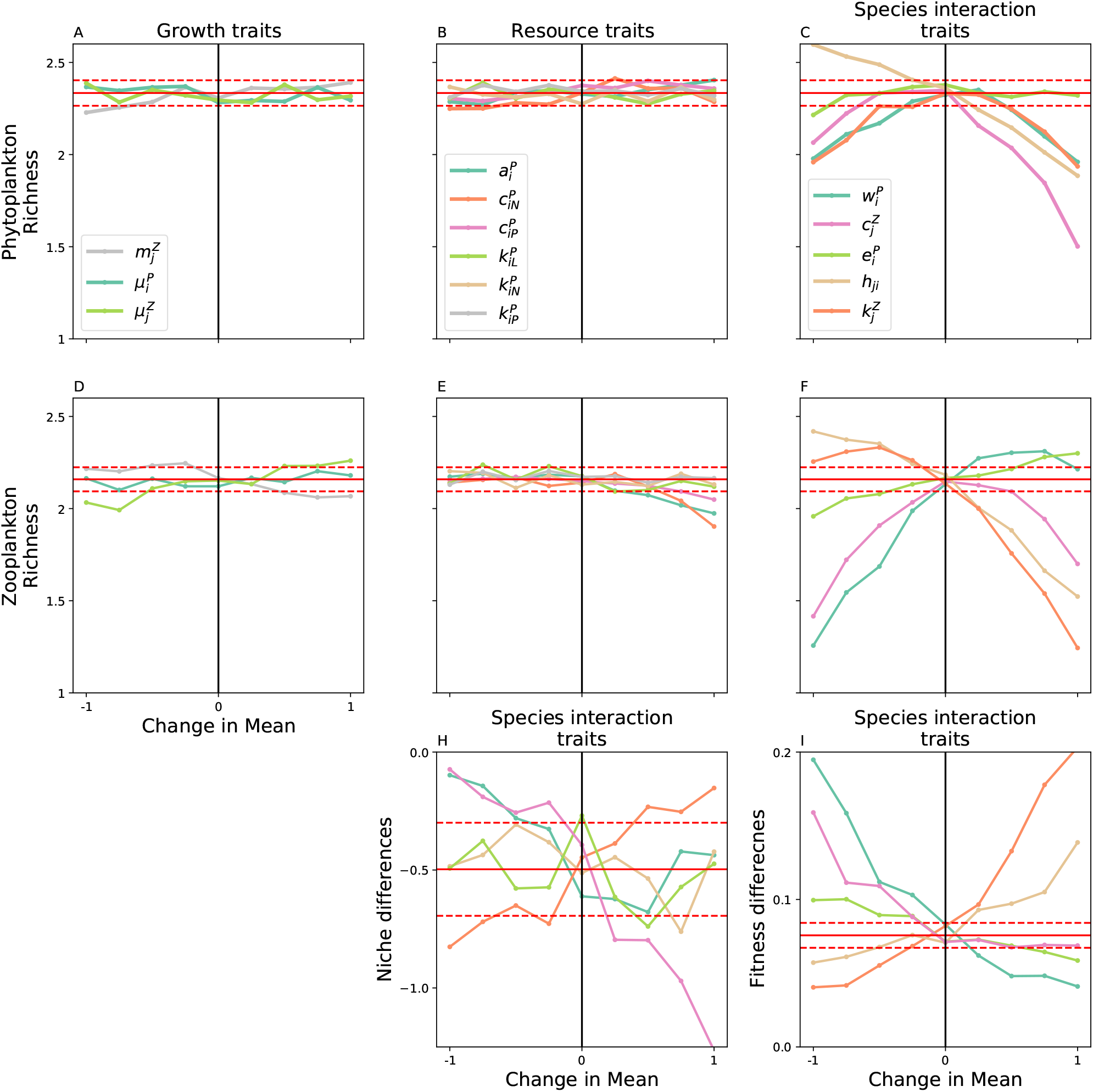
Panels A-F are identical to Figure 2 and here for better comparison. To have a better understanding of the underlying causes for changes in species richness we computed niche and fitness differences for the species interaction traits (Panels C and F). I: Fitness differences explain all changes in species richness that are linked to a decreased viability of zooplankton. H: Conversely, the trait changes do not seem to affect niche differences strongly. The most notable exception is how changes in zooplankton consumption rates 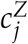 affect niche differences, this explains the non-monotonic dependence of species richness on 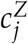. A-I: Black vertical line indicates where all parameters are as they are found empirically. Red horizontal line show the mean and 99% confidence interval of the respective values for the communities generated with empirical values.

**Figure S4:**
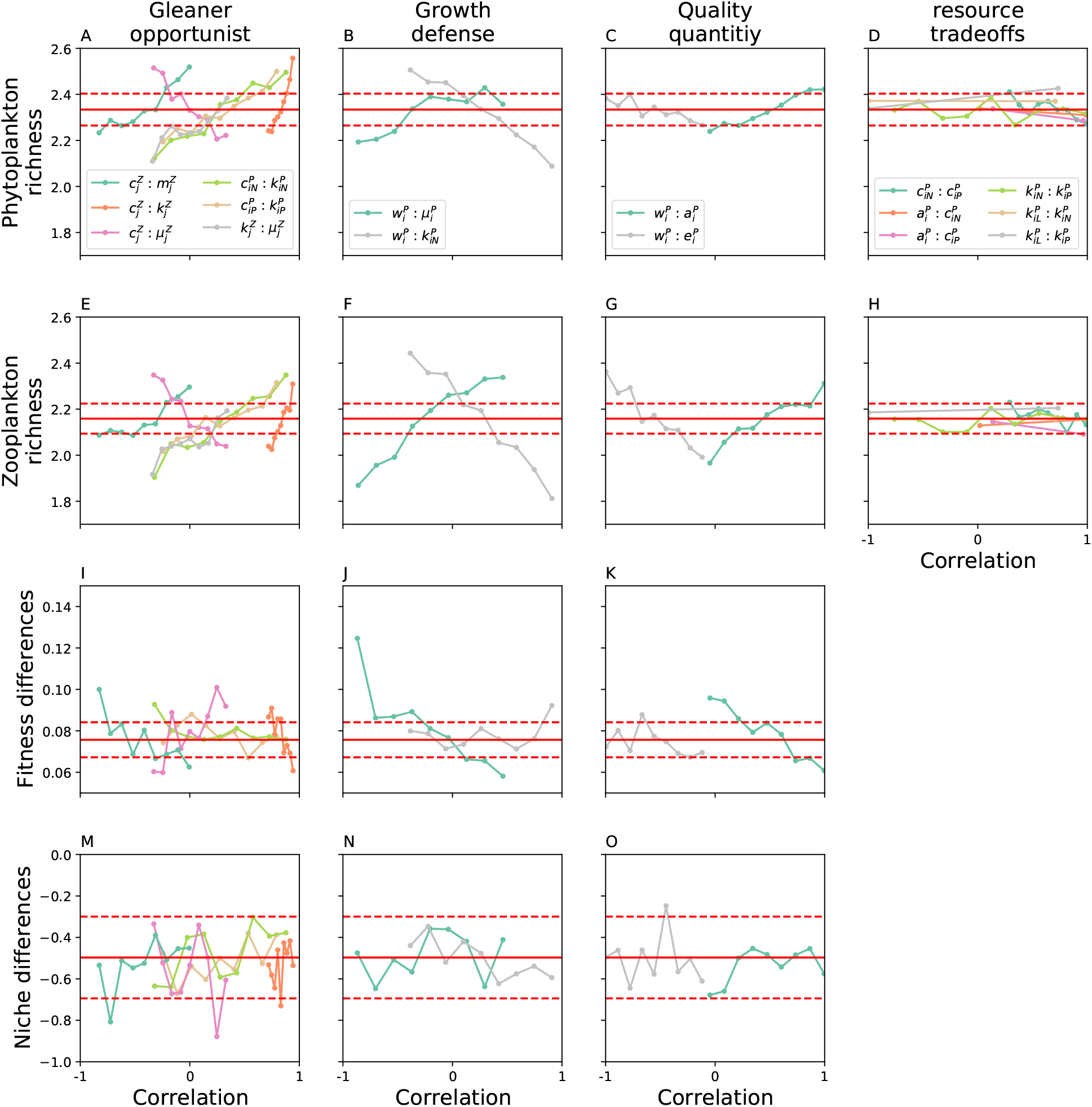
Panels A-H are identical to Figure 3 and here for better comparison. To have a better understanding of the underlying causes for changes in species richness we computed niche and fitness differences for the trait correlations that affected species richness (Panels A-C and E-G). I-K: As for the changes in mean trait values, changes in trade-offs affect fitness differences more strongly than niche differences, which can explain the changes in species richness. M-O: However, the effect on trade-offs on fitness differences is less strong than the effect of changes in mean on fitness differences. A-O: Red horizontal line show the mean and 99% confidence interval of the respective values for the communities generated with empirical values. The correlation cannot be chosen freely for each individual trade-off, rather the maximal and minimal values are determined by the entire trait correlation matrix (see methods).

**Figure S5:**
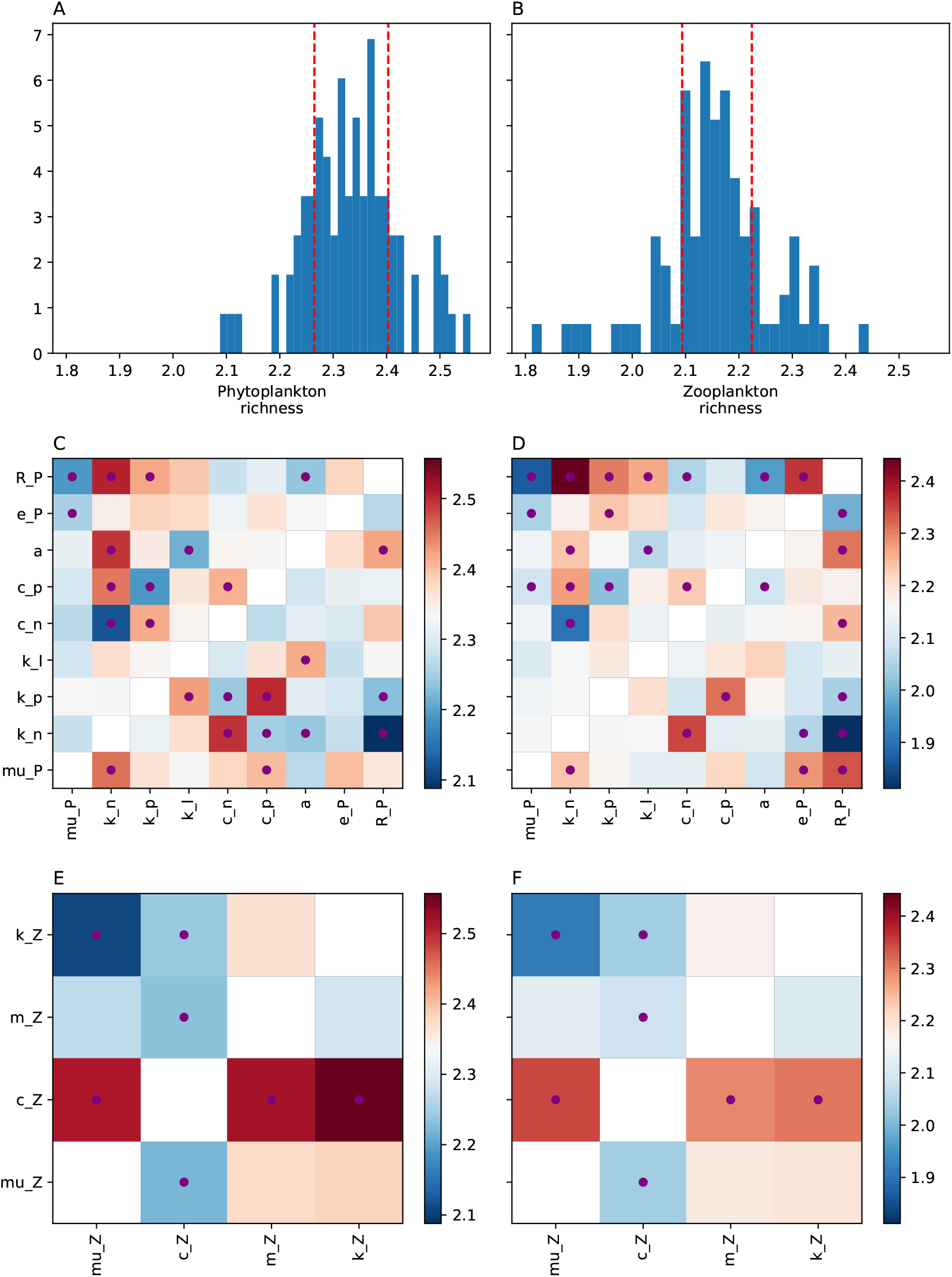
We altered the trait correlation of the phytoplankton and zooplankton trait distributions and assessed their effect on phytoplankton (A,C,E) and zooplankton richness (B,D,F). A,B: Histograms of species richness for phytoplankton (A) and zooplankton (B) for all altered trait distributions. Red dashed lines correspond to the 1% and 99% percentiles of species richness for the reference cases. In the main text we analyzed the ten trait-tradeoffs that deviated most from the reference case. C,D: Color indicates the species richness of phytoplankton (C) and zooplankton (D) for changes in phytoplankton trait distributions (x and y axis). Above diagonal correspond to increased trait-correlations, below diagonal correspond to reduced trait-correlations. Red dots indicate species richness above the 99% or below the 1% species richness for the reference case. E,F Similar to C and D but with altered zooplankton trait distributions.

**Figure S6:**
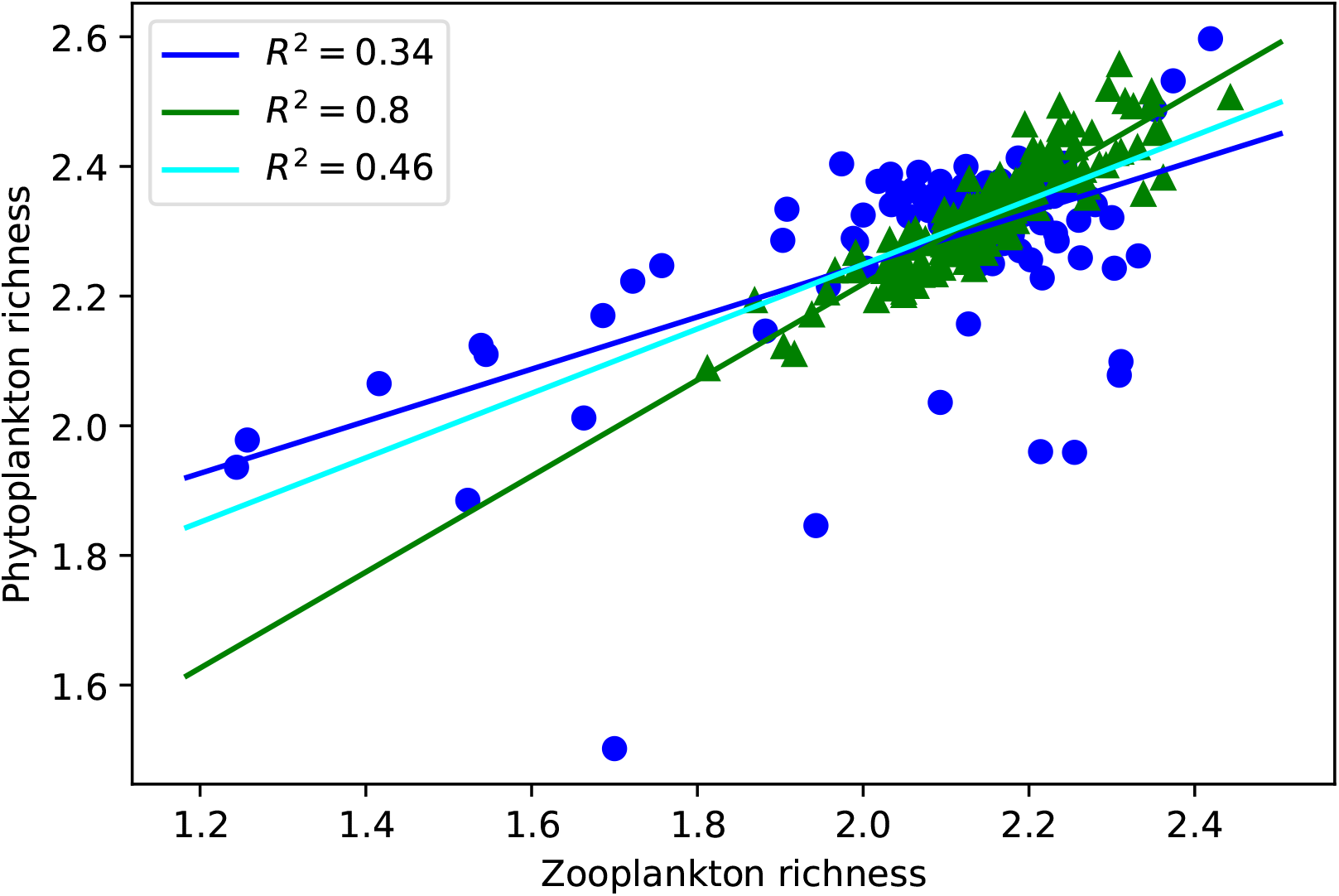
Bi-variate plot of phytoplankton (y-axis) and zooplankton (x-axis) richness in the different simulations with altered mean (blue dots) and altered correlations (green triangles). Legend shows the *R*^2^ value for the corresponding linear regressions, cyan line shows the linear regression for all simulations combined.

**Figure S7:**
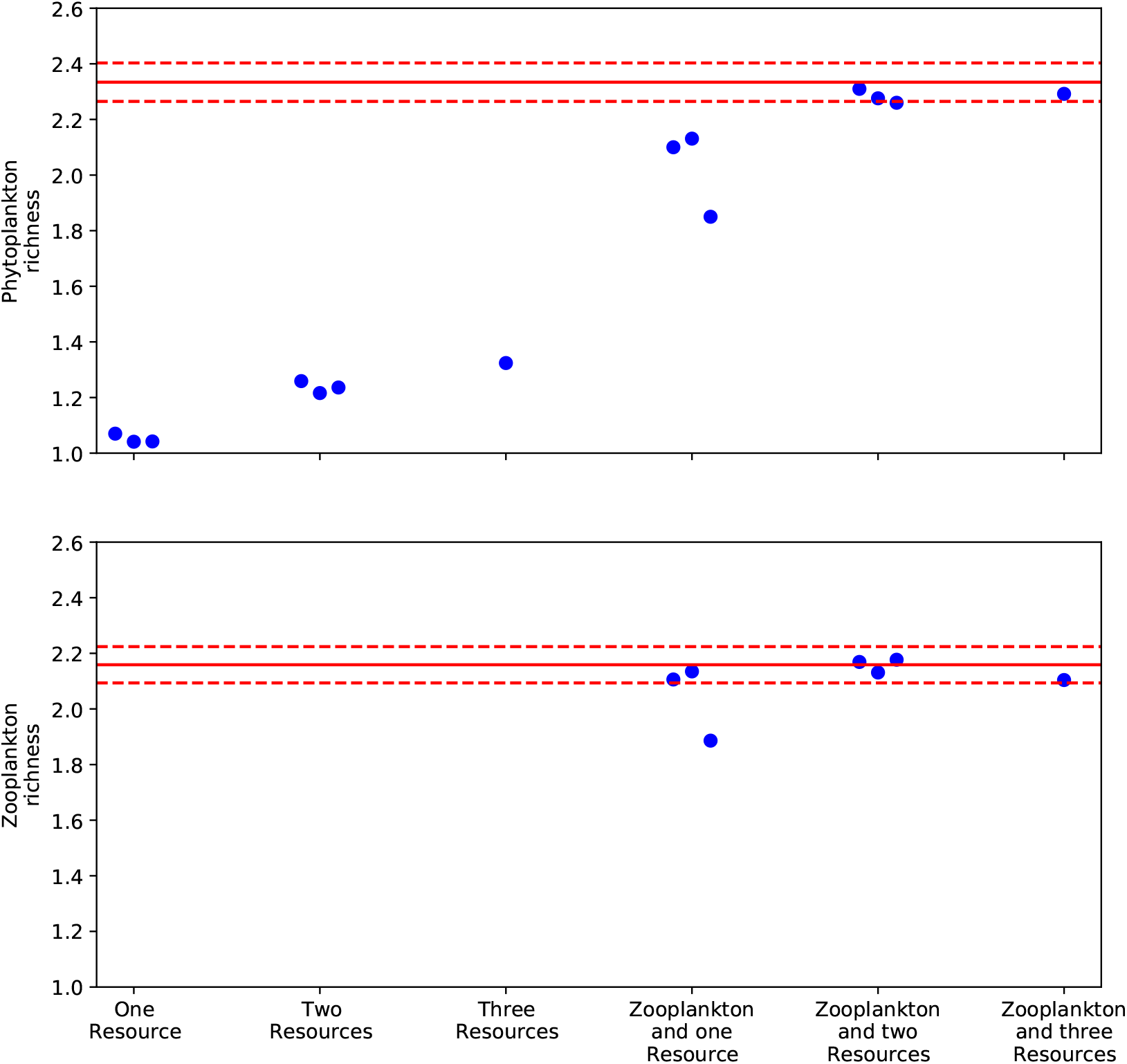
The effect of different mechanisms on species richness. Competition for only one resource (in the absence of predators) leads to the survival of essentially only one species, as predicted by theory. The values are slightly higher because competitive exclusion takes time for competitive similar species. Competition for two or three resources increases species richness, this effect is traditionally reported when investigating the positive effect of resource competition on species coexistence. However, in the presence of zooplankton species the beneficial effect of resource competition is very small. We therefore claim that resource competition is not important for species richness in our community model.

**Figure S8:**
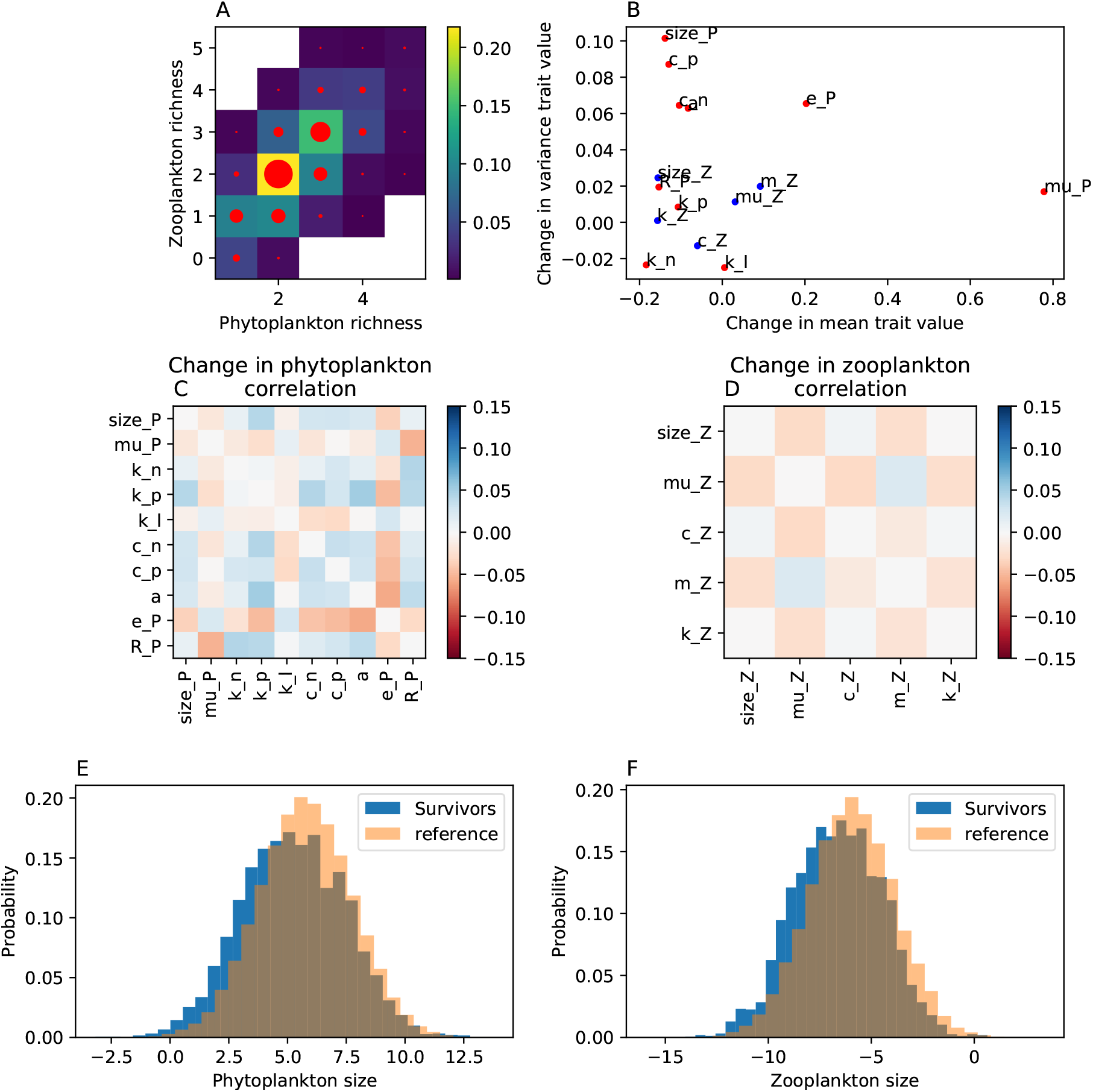
A: 2d-histogram of surviving species. Phytoplankton and zooplankton are well correlated, potentially because zooplankton richness causes niche differentiation for phytoplankton. B: We analyzed the traits of the surviving species and computed mean and variance of each trait. We show the relative difference between trait variance of surviving species and all species (y-axis), and the difference between the trait means, scaled by the variance. In general, larger species tend to survive (Panel E and F). C, D: We computed the trait correlation for the phytoplankton and zooplankton traits of the surviving species. We compared this correlation to the correlation from the meta community. E,F: Histogram of size distribution of the surviving (blue) phytoplankton (E) and zooplankton (F), compared to the size distributions of the meta community (orange). Larger species tend to have a slight fitness advantage.

#### Trait distributions

**Figure S9:**
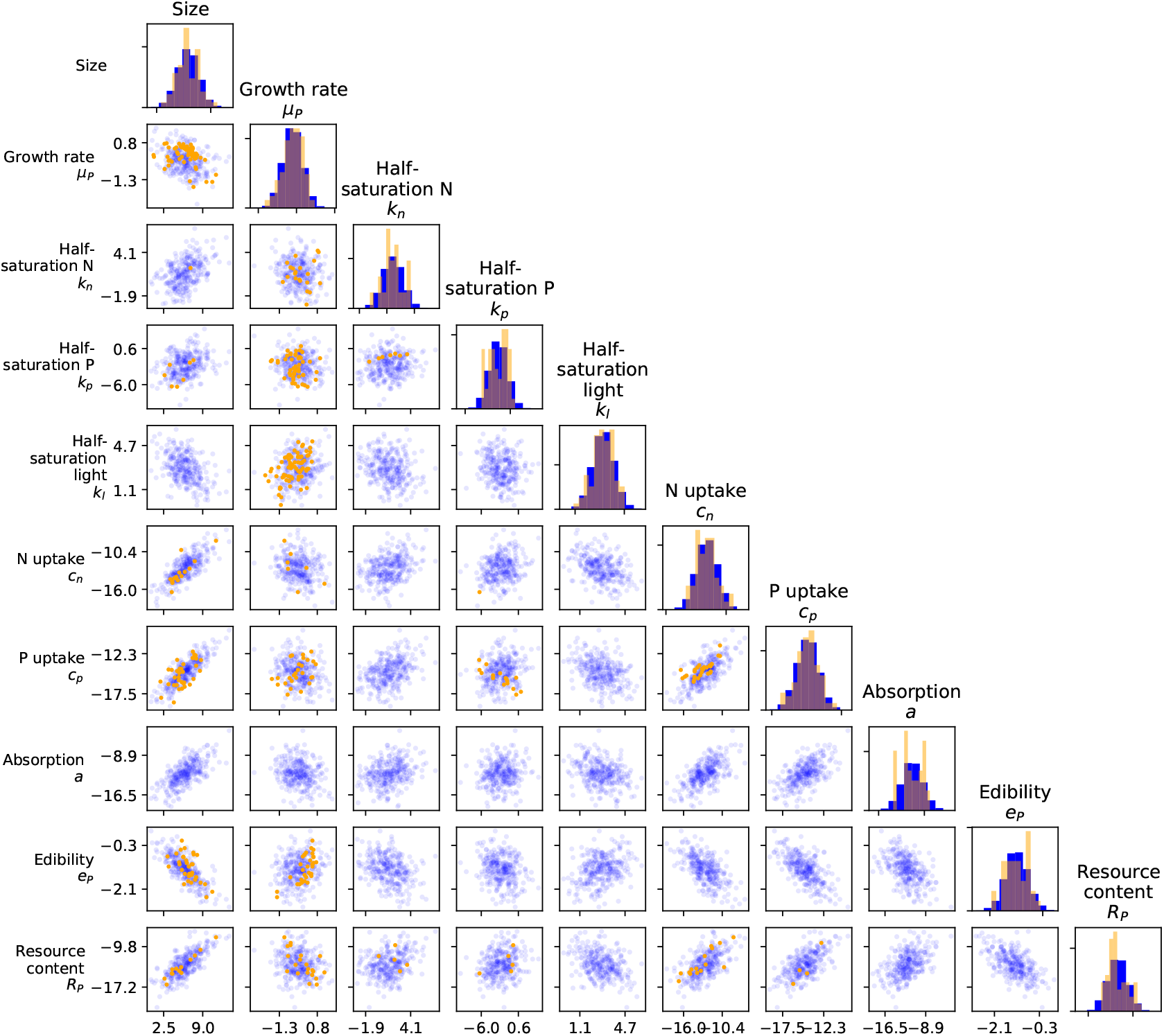
Distribution of phytoplankton traits (diagonal) and their trade-offs (off-diagonal). Diagonal: For each trait we fit a log-normal distribution (blue) to the empirically measured traits values (orange). Off-diagonal: Given these log-normal distribution we use allometric scaling to identify the trade-offs between the traits. In general, this method reproduces the empirically measured phytoplankton species (orange dots) well.

**Figure S10:**
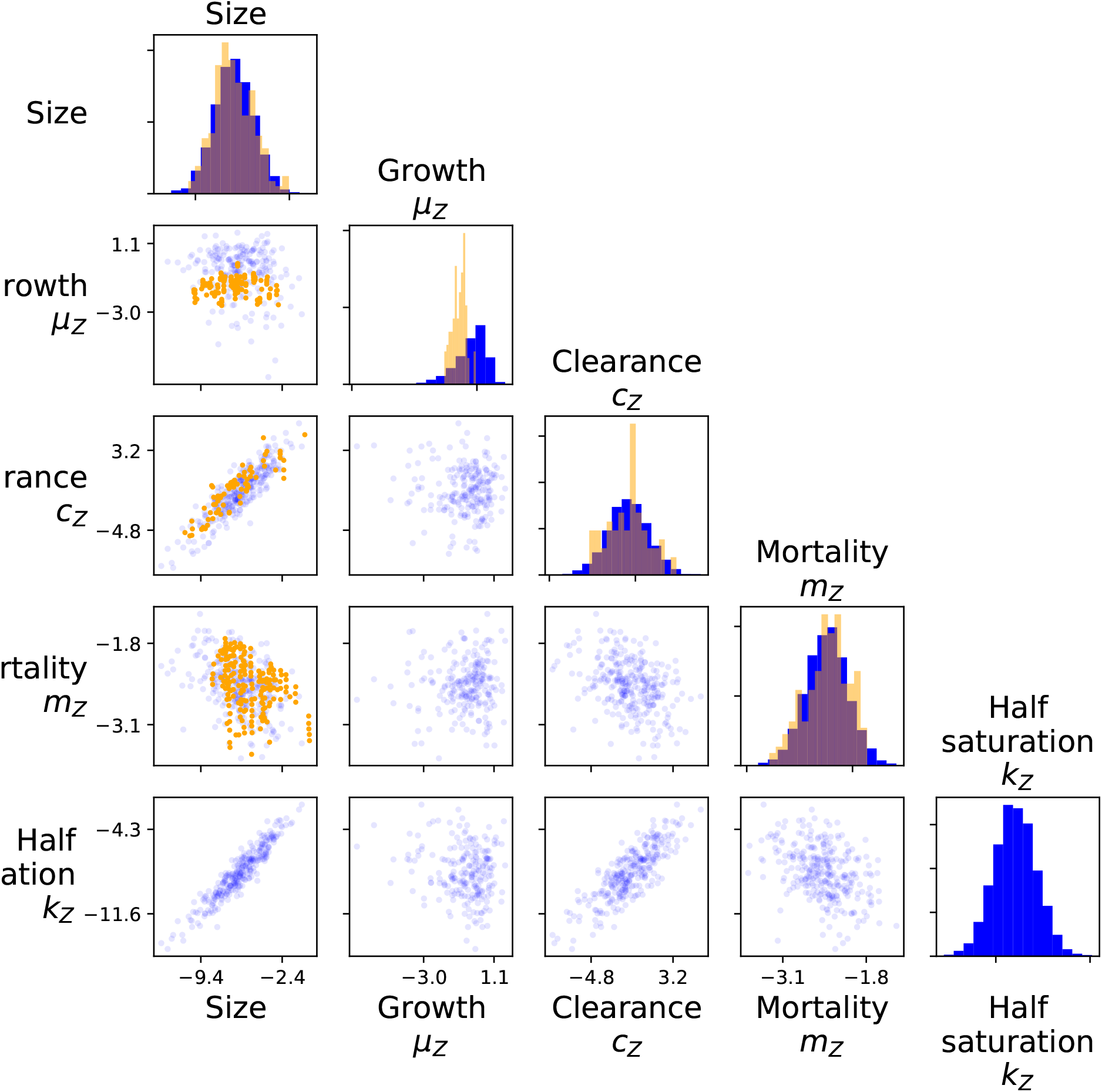
Distribution of zooplankton traits (diagonal) and their trade-offs (off-diagonal). Diagonal: For each trait we fit a log-normal distribution (blue) to the empirically measured traits values (orange). Off-diagonal: Given these log-normal distribution we use allometric scaling to identify the trade-offs between the traits. In general, this method reproduces the empirically measured phytoplankton species (orange dots) well. We did not find any empirically measured values for *k*_*Z*_, but we used the fact that *k*_*Z*_ = *μ*_*Z*_*q*_*Z*_, where *q*_*Z*_ is the resource concentration of the zooplankton, for which empirically measured values are available.

**Figure S11:**
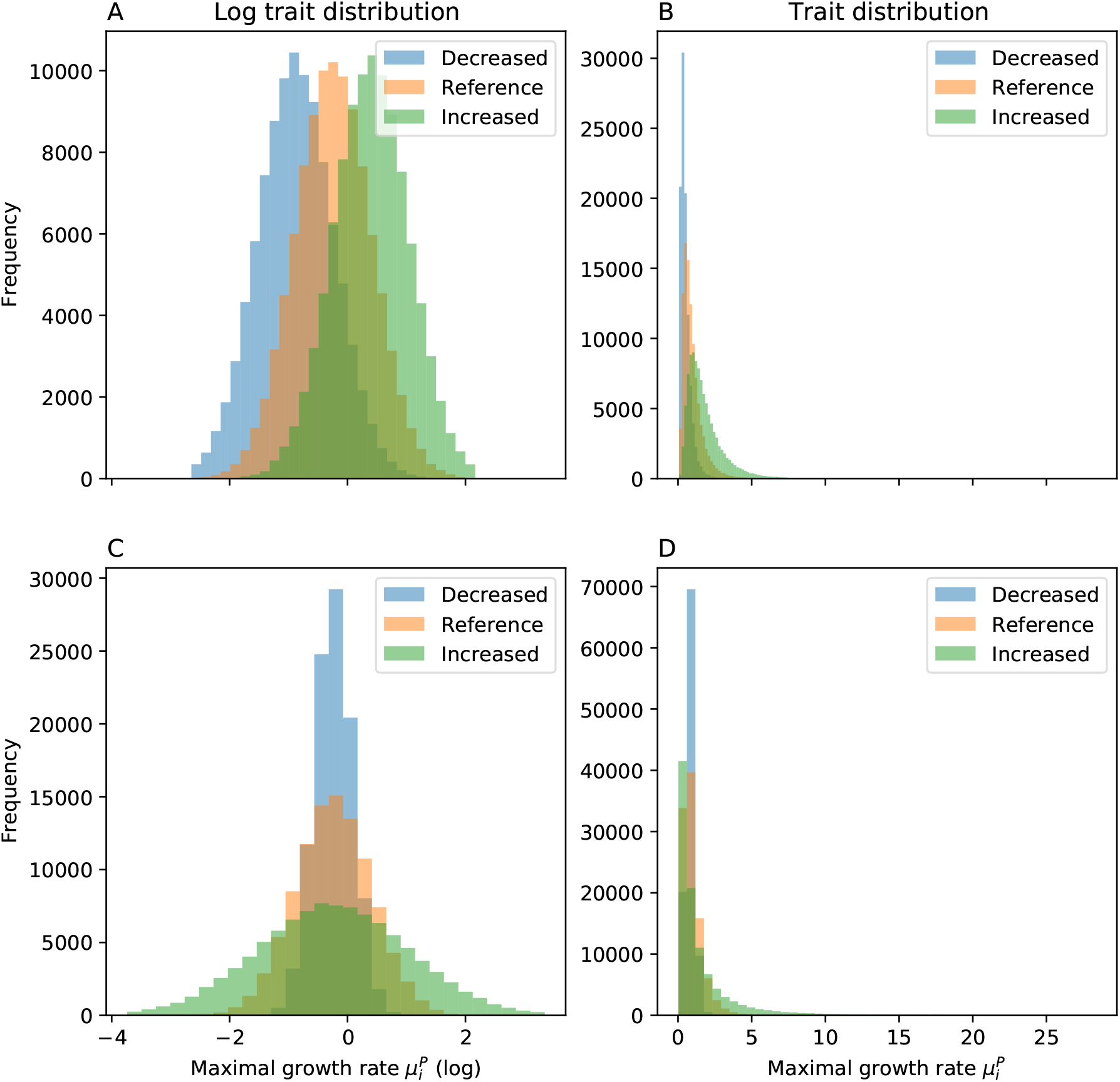
Example of how we altered the trait distribution (here *μ*_*P*_) to assess the importance of the species traits for coexistence. A: The reference distribution (orange) is log-normal, we increased (green) or decreased (blue) the mean of the trait distributions up to one standard deviation. B: Similarly, we increased or decreased the variance by a factor of two to assess the importance of the variance on species coexistence. C, D: Equal to A and B, but shown are the non-log distributions.

### S2 Droop equation

To derive the zooplankton growth equations we assume that zooplankton growth is governed by internal nutrient contents, also known as the Droop equations (Droop, 1973).

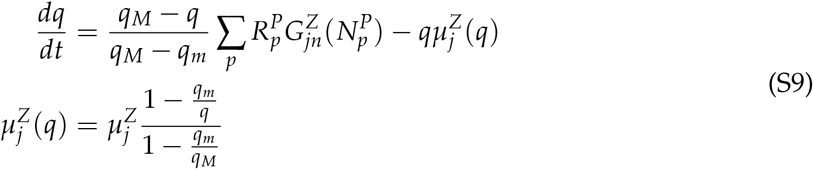

where *q* is the internal resource concentration, *q*_*m*_ is the minimal internal resource concentration, *q*_*M*_ is the maximal internal resource concentration. 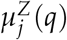 is the growth rate of the zooplankton as a function of the internal resource concentration *q*, note that growth at maximal internal resource concentration equals the maximum growth rate of the zooplankton, i.e.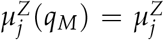. We assume that the internal resource concentration is always at equilibrium, i.e. 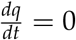, which leads to

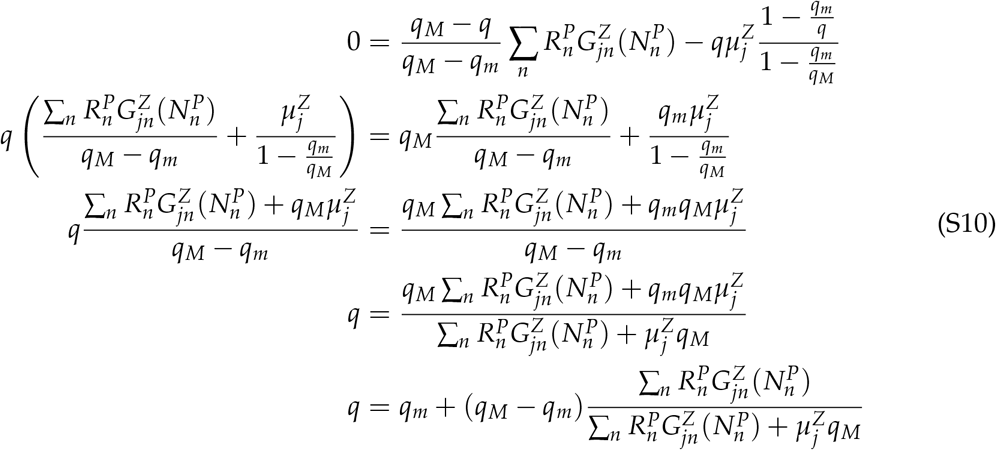

Inserting this into the growth rate 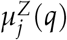 yields:

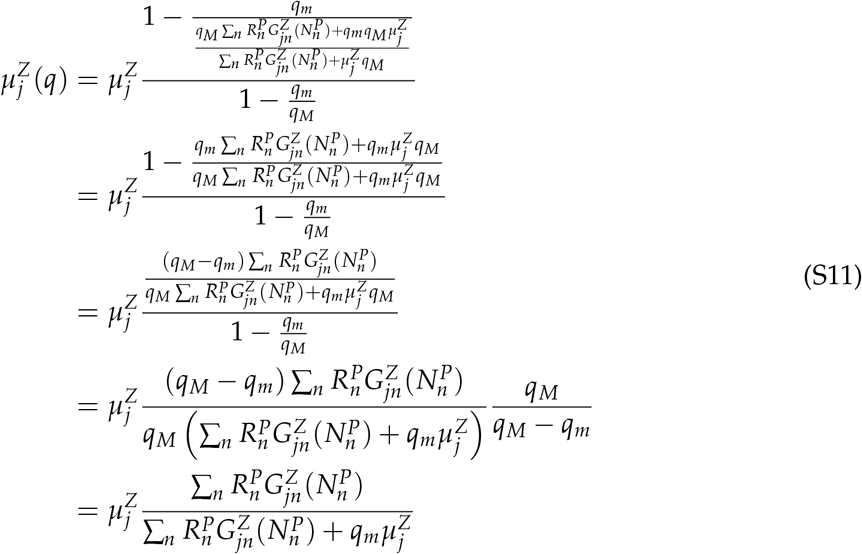

We therefore find that 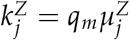.

#### Phytoplankton

To derive the phytoplankton growth equations we assume that phytoplankton growth is governed by internal nutrient contents, also known as the Droop equations.

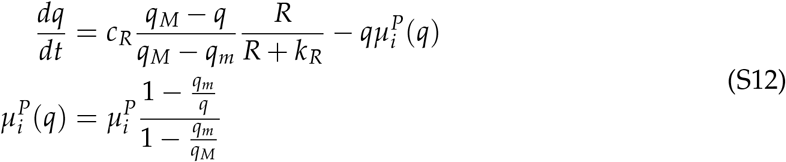

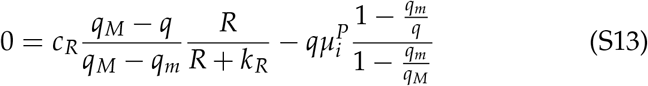

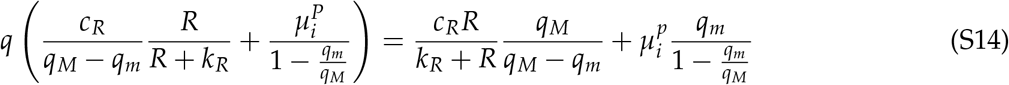

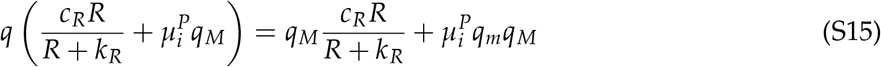

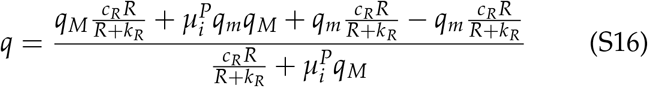

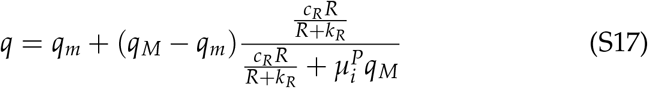

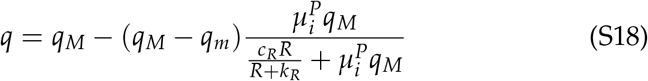

We insert S17 into S12:

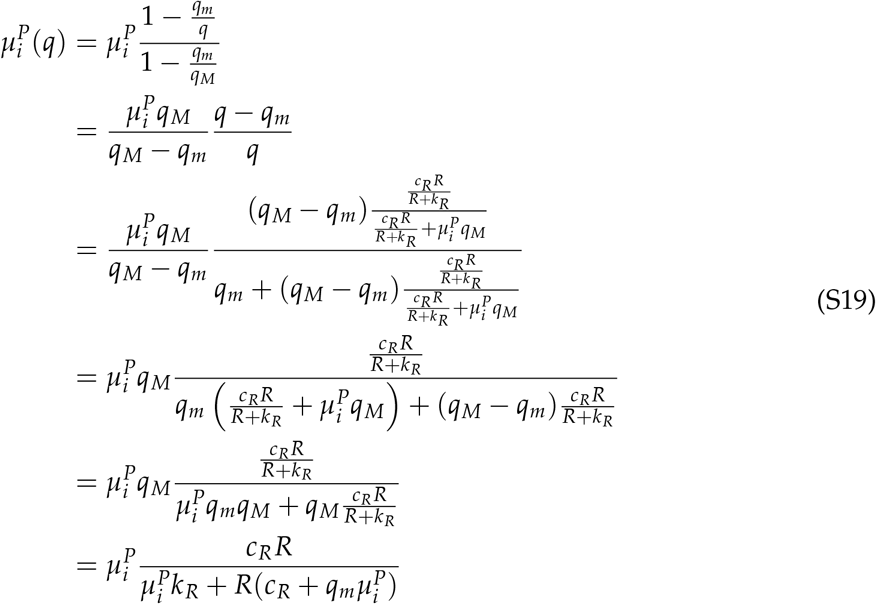

Similarly, we can compute the resource uptake dynamics of the phytoplankton

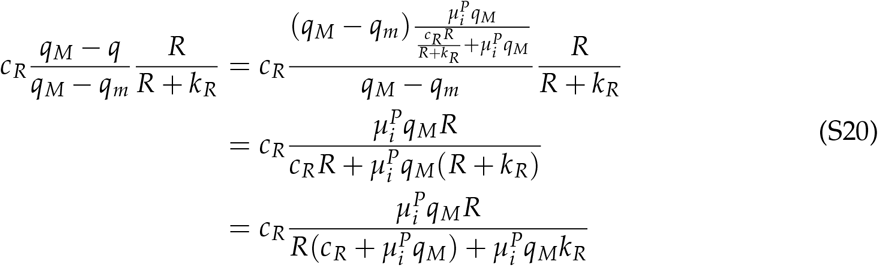

#### Units verification

Here we verify that the model equations are dimensionally correct.

Resource consumption:

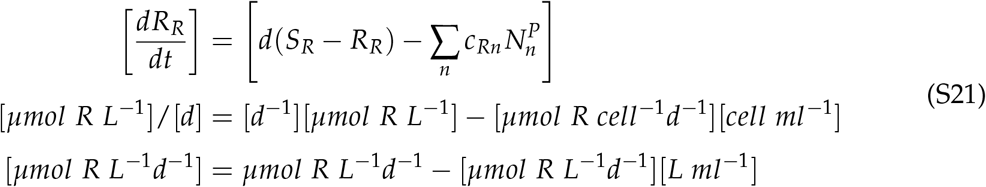

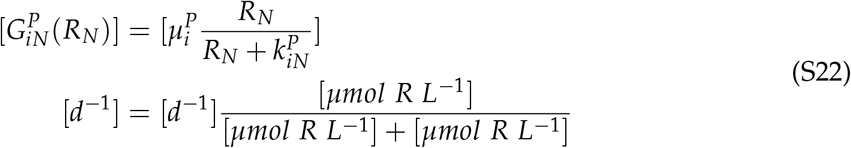

For light competition we first compute

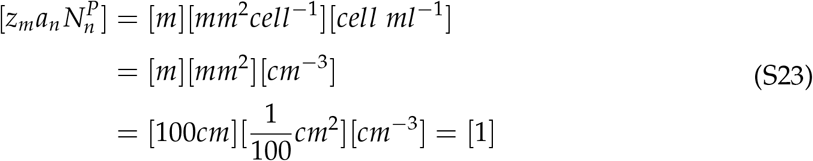

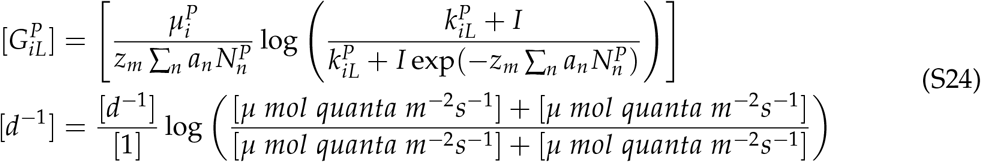

Grazing of phytoplankton

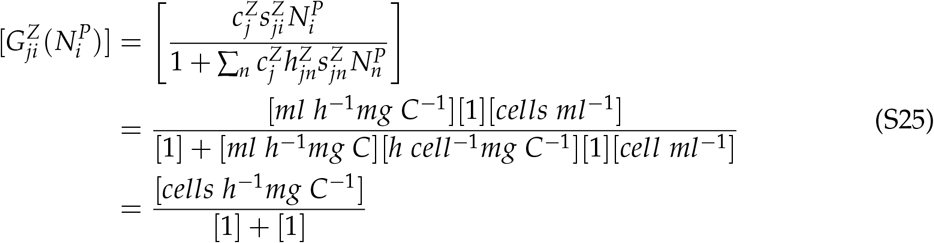

Phytoplankton growth rate

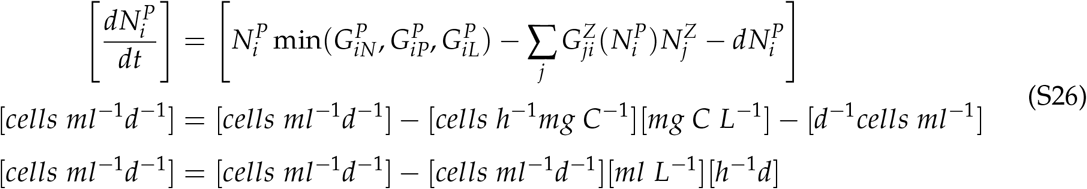

Zooplankton growth rate

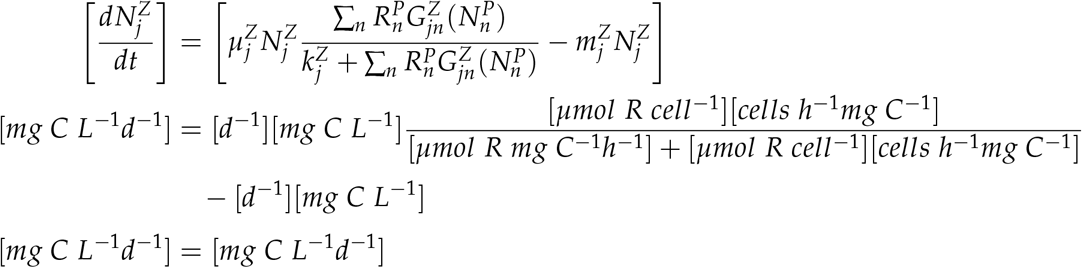

## References

Adler, P.B., Ellner, S.P. & Levine, J.M. (2010). Coexistence of perennial plants: an embarrassment of niches: Embarrassment of niches. Ecology Letters, pp. no–no.

Angert, A.L., Huxman, T.E., Chesson, P. & Venable, D.L. (2009). Functional tradeoffs determine species coexistence via the storage effect. Proceedings of the National Academy of Sciences, 106, 11641–11645.

Armstrong, R. & McGehee, R. (1980). Competitive exclusion. The American Naturalist, 115, 151–170.

Armstrong, R.A. & McGehee, R. (1976). Coexistence of species competing for shared resources. Theoretical Population Biology, 9, 317–328.

Arsenault, R. & Owen-Smith, N. (2002). Facilitation versus competition in grazing herbivore assemblages. Oikos, 97, 313–318.

Bagchi, R., Gallery, R.E., Gripenberg, S., Gurr, S.J., Narayan, L., Addis, C.E., Freckleton, R.P. & Lewis, O.T. (2014). Pathogens and insect herbivores drive rainforest plant diversity and composition. Nature, 506, 85–88.

Barabás, G., D’Andrea, R. & Stump, S.M. (2018). Chesson’s coexistence theory. Ecological Monographs, 88, 277–303.

Becerra, J.X. (2015). On the factors that promote the diversity of herbivorous insects and plants in tropical forests. Proceedings of the National Academy of Sciences, 112, 6098–6103.

Berggreen, U., Hansen, B. & Kiørboe, T. (1988). Food size spectra, ingestion and growth of the copepodAcartia tonsa during development: Implications for determination of copepod production. Marine Biology, 99, 341–352.

Branco, P., Egas, M., Hall, S.R. & Huisman, J. (2020). Why Do Phytoplankton Evolve Large Size in Response to Grazing? The American Naturalist, 195, E20–E37.

Brun, P., Payne, M.R. & Kiørboe, T. (2016). A trait database for marine copepods, supplement to: Brun, Philipp; Payne, Mark R. Kiørboe, Thomas (2017): A trait database for marine copepods. Earth System Science Data, 9(1), 99-113. Artwork Size: 1.2 MBytes Medium: application/octet-stream Pages: 1.2 MBytes type: dataset.

Buche, L., Spaak, J.W., Jarillo, J. & De Laender, F. (2021). Niche difference determines coexistence and similar underlying processes in four ecological groups. preprint, Ecology.

Carroll, I.T., Cardinale, B.J. & Nisbet, R.M. (2011). Niche and fitness differences relate the maintenance of diversity to ecosystem function. 92, 9.

Chesson, P. (1990). MacArthur’s consumer-resource model. Theoretical Population Biology, 37, 26–38.

Chesson, P. (1994). Multispecies Competition in Variable Environments. Theoretical Population Biology, 45, 227–276.

Chesson, P. (2000). Mechanisms of Maintenance of Species Diversity. Annual Review of Ecology and Systematics, 31, 343–366.

Chesson, P. (2003). Quantifying and testing coexistence mechanisms arising from recruitment fluctuations. Theoretical Population Biology, 64, 345–357.

Chesson, P. (2018). Updates on mechanisms of maintenance of species diversity. Journal of Ecology, 106, 1773–1794.

Chesson, P. & Kuang, J.J. (2008). The interaction between predation and competition. Nature, 456, 235–238.

Creed, R.P. (2000). Is there a new keystone species in North American lakes and rivers? Oikos, 91, 405–408.

Droop, M.R. (1973). SOME THOUGHTS ON NUTRIENT LIMITATION IN ALGAE1. Journal of Phycology, 9, 264–272. eprint: https://onlinelibrary-wiley-com.proxy.library.cornell.edu/doi/pdf/10.1111/j.1529-8817.1973.tb04092.x.

Edwards, K.F., Klausmeier, C.A. & Litchman, E. (2011). Evidence for a three-way trade-off between nitrogen and phosphorus competitive abilities and cell size in phytoplankton. Ecology, 92, 2085–2095.

Edwards, K.F., Thomas, M.K., Klausmeier, C.A. & Litchman, E. (2012). Allometric scaling and taxonomic variation in nutrient utilization traits and maximum growth rate of phytoplankton. Limnology and Oceanography, 57, 554–566.

Ehrlich, E., Kath, N.J. & Gaedke, U. (2020). The shape of a defense-growth trade-off governs seasonal trait dynamics in natural phytoplankton. The ISME Journal, 14, 1451–1462.

Ellner, S.P., Snyder, R.E., Adler, P.B. & Hooker, G. (2019). An expanded modern coexistence theory for empirical applications. Ecology Letters, 22, 3–18.

Field, C.B. (1998). Primary Production of the Biosphere: Integrating Terrestrial and Oceanic Components. Science, 281, 237–240.

Finkel, Z.V., Beardall, J., Flynn, K.J., Quigg, A., Rees, T.A.V. & Raven, J.A. (2010). Phytoplankton in a changing world: cell size and elemental stoichiometry. Journal of Plankton Research, 32, 119–137.

Gallego, I., Venail, P. & Ibelings, B.W. (2019). Size differences predict niche and relative fitness differences between phytoplankton species but not their coexistence. The ISME Journal, 13, 1133–1143.

Germain, R.M., Weir, J.T. & Gilbert, B. (2016). Species coexistence: macroevolutionary relationships and the contingency of historical interactions. Proceedings of the Royal Society B: Biological Sciences, 283, 20160047.

Godoy, O., Bartomeus, I., Rohr, R.P. & Saavedra, S. (2018). Towards the Integration of Niche and Network Theories. Trends in Ecology & Evolution, 33, 287–300.

Godoy, O. & Levine, J.M. (2014). Phenology effects on invasion success: insights from coupling field experiments to coexistence theory. Ecology, 95, 726–736.

Huisman, J. & Olff, H. (1998). Competition and facilitation in multispecies plant-herbivore systems of productive environments. Ecology Letters, 1998, 25–29.

Huisman, J., Pham Thi, N.N., Karl, D.M. & Sommeijer, B. (2006). Reduced mixing generates oscillations and chaos in the oceanic deep chlorophyll maximum. Nature, 439, 322–325.

Huisman, J. & Weissing, F.J. (1994). Light-Limited Growth and Competition for Light in Well-Mixed Aquatic Environments: An Elementary Model. Ecology, 75, 507–520.

Huisman, J. & Weissing, F.J. (1999). Biodiversity of plankton by species oscillations and chaos. Nature, 402, 407–410.

Hutchinson, G.E. (1959). Homage to Santa Rosalia or Why Are There So Many Kinds of Animals? The American Naturalist, 93, 145–159.

Janzen, D.H. (1970). Herbivores and the Number of Tree Species in Tropical Forests. The American Naturalist, 104, 501–528.

Kandlikar, G.S., Johnson, C.A., Yan, X., Kraft, N.J.B. & Levine, J.M. (2019). Winning and losing with microbes: how microbially mediated fitness differences influence plant diversity. Ecology Letters, p. ele.13280.

Kiørboe, T., Visser, A. & Andersen, K.H. (2018). A trait-based approach to ocean ecology. ICES Journal of Marine Science, 75, 1849–1863.

Kraft, N.J.B., Godoy, O. & Levine, J.M. (2015). Plant functional traits and the multidimensional nature of species coexistence. Proceedings of the National Academy of Sciences, 112, 797–802.

Letten, A.D., Dhami, M.K., Ke, P.J. & Fukami, T. (2018). Species coexistence through simultaneous fluctuation-dependent mechanisms. Proceedings of the National Academy of Sciences, 115, 6745–6750.

Letten, A.D., Ke, P.J. & Fukami, T. (2017). Linking modern coexistence theory and contemporary niche theory. Ecological Monographs, 87, 161–177.

Letten, A.D. & Stouffer, D.B. (2019). The mechanistic basis for higher-order interactions and non-additivity in competitive communities. Ecology Letters, 22, 423–436.

Levine, J.M. & HilleRisLambers, J. (2009). The importance of niches for the maintenance of species diversity. Nature, 461, 254–257.

Lind, E.M., Borer, E.T., Seabloom, E., Adler, P.B., Bakker, J.D., Blumenthal, D.M., Crawley, M., Davies, K., Firn, J., Gruner, D.S., Harpole, W.S., Hautier, Y., Hillebrand, H., Knops, J.M.H., Melbourne, B.A., Mortensen, B., Risch, A.C., Schuetz, M., Stevens, C. & Wragg, P.D. (2013). Life-history constraints in grassland plant species: a growth-defence trade-off is the norm. Ecology Letters, 2019, 513–521.

Litchman, E. (2003). Competition and coexistence of phytoplankton under fluctuating light: experiments with two cyanobacteria. Aquatic Microbial Ecology, 31, 241–248.

Litchman, E. & Klausmeier, C.A. (2001). Competition of Phytoplankton under Fluctuating Light. p. 18.

Litchman, E. & Klausmeier, C.A. (2008). Trait-Based Community Ecology of Phytoplankton. Annual Review of Ecology, Evolution, and Systematics, 39, 615–639.

Litchman, E., Klausmeier, C.A., Schofield, O.M. & Falkowski, P.G. (2007). The role of functional traits and trade-offs in structuring phytoplankton communities: scaling from cellular to ecosystem level. Ecology Letters, 10, 1170–1181.

Litchman, E., Ohman, M.D. & Kiørboe, T. (2013). Trait-based approaches to zooplankton communities. Journal of Plankton Research, 35, 473–484.

Merkli, S. (2021). The relative importance of top-down and bottom-up controls on the growth rate of phytoplankton trait-based groups. Master’s thesis, University Zurich.

Narwani, A., Alexandrou, M.A., Oakley, T.H., Carroll, I.T. & Cardinale, B.J. (2013). Experimental evidence that evolutionary relatedness does not affect the ecological mechanisms of coexistence in freshwater green algae. Ecology Letters, 16, 1373–1381.

Narwani, A., Bentlage, B., Alexandrou, M.A., Fritschie, K.J., Delwiche, C., Oakley, T.H. & Cardinale, B.J. (2017). Ecological interactions and coexistence are predicted by gene expression similarity in freshwater green algae. Journal of Ecology, 105, 580–591.

Olff, H. & Ritchie, M.E. (1998). Effects of herbivores on grassland plant diversity. Trends in Ecology & Evolution, 13, 261–265.

Pande, J., Fung, T., Chisholm, R. & Shnerb, N.M. (2019). Mean growth rate when rare is not a reliable metric for persistence of species. preprint, Ecology.

Pérez-Ramos, I.M., Matías, L., Gómez-Aparicio, L. & Godoy, O. (2019). Functional traits and phenotypic plasticity modulate species coexistence across contrasting climatic conditions. Nature Communications, 10, 2555.

Schreiber, S.J. (2000). Criteria for Cr Robust Permanence. Journal of Differential Equations, 162, 400–426.

Shoemaker, L.G., Barner, A.K., Bittleston, L.S. & Teufel, A.I. (2019). Quantifying the relative importance of competition, predation, and environmental variation for species coexistence. preprint, Ecology.

Shoemaker, L.G., Barner, A.K., Bittleston, L.S. & Teufel, A.I. (2020a). Quantifying the relative importance of variation in predation and the environment for species coexistence. Ecology Letters, 23, 939–950.

Shoemaker, L.G., Sullivan, L.L., Donohue, I., Cabral, J.S., Williams, R.J., Mayfield, M.M., Chase, J.M., Chu, C., Harpole, W.S., Huth, A., HilleRisLambers, J., James, A.R.M., Kraft, N.J.B., May, F., Muthukrishnan, R., Satterlee, S., Taubert, F., Wang, X., Wiegand, T., Yang, Q. & Abbott, K.C. (2020b). Integrating the underlying structure of stochasticity into community ecology. Ecology, 101.

Singh, P. & Baruah, G. (2020). Higher order interactions and species coexistence. Theoretical Ecology.

Spaak, J., Millet, R., Ke, P.J., Letten, A.D. & De Laender, F. (2021a). The effect of non-linear competitive interactions on quantifying niche and fitness differences. preprint, Ecology.

Spaak, J.W., Carpentier, C. & De Laender, F. (2021b). Species richness increases fitness differences, but does not affect niche differences. Ecology Letters, p. ele.13877.

Spaak, J.W. & De Laender, F. (2020). Intuitive and broadly applicable definitions of niche and fitness differences. Ecology Letters, 23, 1117–1128.

Spaak, J.W. & De Laender, F. (2021). Effects of pigment richness and size variation on coexistence, richness and function in light-limited phytoplankton. Journal of Ecology, pp. 1365–2745.13645.

Spaak, J.W., Godoy, O. & De Laender, F. (2021c). Mapping species niche and fitness differences for communities with multiple interaction types. Oikos, p. oik.08362.

Stomp, M., Huisman, J., de Jongh, F., Veraart, A.J., Gerla, D., Rijkeboer, M., Ibelings, B.W., Wollenzien, U.I.A. & Stal, L.J. (2004). Adaptive divergence in pigment composition promotes phytoplankton biodiversity. Nature, 432, 104–107.

Tilman, D., Kilham, S.S. & Kilham, P. (1982). Phytoplankton Community Ecology: The Role of Limiting Nutrients. Annual Review of Ecology and Systematics, 13, 349–372.

Usinowicz, J., Chang-Yang, C.H., Chen, Y.Y., Clark, J.S., Fletcher, C., Garwood, N.C., Hao, Z., Johnstone, J., Lin, Y., Metz, M.R., Masaki, T., Nakashizuka, T., Sun, I.F., Valencia, R., Wang, Y., Zimmerman, J.K., Ives, A.R. & Wright, S.J. (2017). Temporal coexistence mechanisms contribute to the latitudinal gradient in forest diversity. Nature, 550, 105–108.

Usinowicz, J., Wright, S.J. & Ives, A.R. (2012). Coexistence in tropical forests through asynchronous variation in annual seed production. Ecology, 93, 2073–2084.

Uye, S.i. (1982). Length-Weight Relationships of Important Zooplankton from the Inland Sea of Japan. Journal of the Oceanographical Society of Japan, 38, 149–158.

Yamamichi, M., Kyogoku, D., Iritani, R., Kobayashi, K., Takahashi, Y., Tsurui-Sato, K., Yamawo, A., Dobata, S., Tsuji, K. & Kondoh, M. (2020). Intraspecific Adaptation Load: A Mechanism for Species Coexistence. Trends in Ecology & Evolution, 35, 897–907.

Zepeda, V. & Martorell, C. (2019). Fluctuation-independent niche differentiation and relative non-linearity drive coexistence in a species-rich grassland. Ecology, 100.

